# Changes in the cellular makeup of motor patterning circuits drive courtship song evolution in *Drosophila*

**DOI:** 10.1101/2024.01.23.576861

**Authors:** Dajia Ye, Justin T. Walsh, Ian P. Junker, Yun Ding

## Abstract

How evolutionary changes in genes and neurons encode species variation in complex motor behaviors are largely unknown. Here, we develop genetic tools that permit a neural circuit comparison between the model species *Drosophila melanogaster* and the closely-related species *D. yakuba*, who has undergone a lineage-specific loss of sine song, one of the two major types of male courtship song in *Drosophila*. Neuroanatomical comparison of song patterning neurons called TN1 across the phylogeny demonstrates a link between the loss of sine song and a reduction both in the number of TN1 neurons and the neurites serving the sine circuit connectivity. Optogenetic activation confirms that TN1 neurons in *D. yakuba* have lost the ability to drive sine song, while maintaining the ability to drive the singing wing posture. Single-cell transcriptomic comparison shows that *D. yakuba* specifically lacks a cell type corresponding to TN1A neurons, the TN1 subtype that is essential for sine song. Genetic and developmental manipulation reveals a functional divergence of the sex determination gene *doublesex* in *D. yakuba* to reduce TN1 number by promoting apoptosis. Our work illustrates the contribution of motor patterning circuits and cell type changes in behavioral evolution, and uncovers the evolutionary lability of sex determination genes to reconfigure the cellular makeup of neural circuits.

## Introduction

Behavioral diversity is widespread and essential for species-specific adaptations in survival and reproduction ^1,2^. Defining how species differences in behaviors are encoded by evolutionary changes in genes and nervous systems is challenging in any system, especially for complex motor behaviors because they involve an intricate interplay among circuit elements that transform sensory inputs to motor outputs. Where and how nervous systems have evolved to generate species diversity in complex motor behaviors remain largely unaddressed. Due to their rich behavioral differences and experimental accessibility, *Drosophila* species recently emerged as a promising system for species comparisons to explore the mechanisms by which behaviors evolve^3^, for example, linking species-specific host preferences and mate recognition to the evolution of sensory receptors and sensory circuits^4–9^.

The rapid diversification of *Drosophila* courtship song and the characterization of genes and circuits in song generation in the model species *D. melanogaster* together provide an opportunity to investigate the evolution of complex motor behaviors^10^. Driven by sexual selection, courtship behaviors are exceptionally diverse, providing some of the most spectacular examples of behavioral diversification and elaboration^11,12^. In *Drosophila* species, males perform a sophisticated courtship ritual that involves a series of motor elements, such as chasing, circling, singing, and licking^13^. Males sing by vibrating their wings in specific motions to generate a species-specific and context-dependent song that females use for mate selection^14,15^. The courtship song of *Drosophila* species, including *D. melanogaster*, comprises two major types: pulse song, trains of discrete pulses, and sine song, continuous sinusoidal waves^14,15^ (**Fig. 1a**). Males continuously adjust the use of song types during the dynamic social interaction of courtship^16^.

**Fig. 1.**
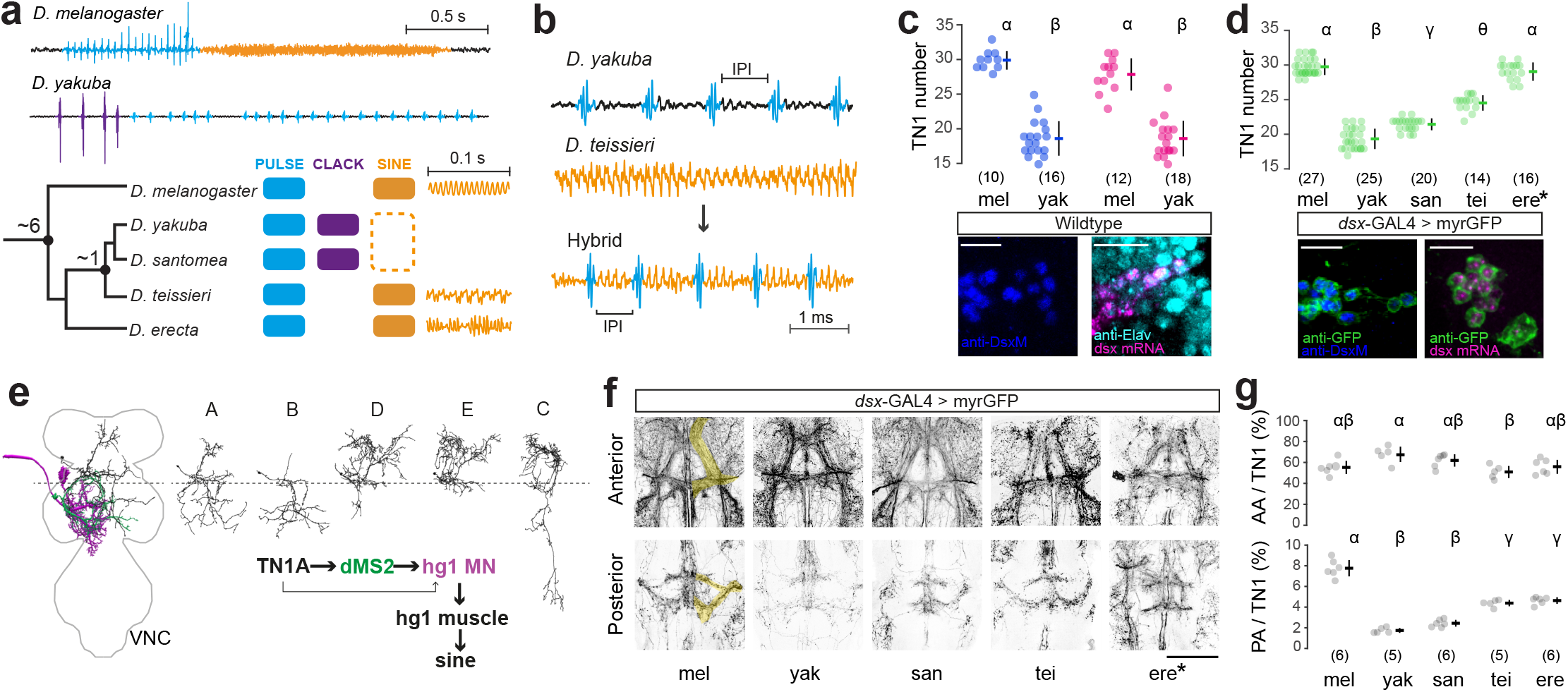
Species diversity of male courtship song and TN1 neuroanatomy. **(a)** Top: representative song traces showing courtship song types in *D. melanogaster* (pulse and sine) and *D. yakuba* (pulse and clack). Bottom: Phylogeny (divergence time noted in million years based on^70,71^) of five species in the *melanogaster* subgroup, with song types and exemplar waveforms of sine song shown on the right. **(b)** F1 hybrid males between *D. yakuba* and *D. teissieri* sing a pulse song (blue) with an IPI resembling *D. teissieri* sine song (orange). Also see **Fig. S1**. **(c)** Cell number of the *dsx*-expressing TN1 neurons (per hemisphere) measured by IHC (blue) and HCR (magenta) in the wildtype strains of *D. melanogaster* (mel) and *D. yakuba* (yak). Representative confocal images are shown on the bottom. **(d)** Cell number measured by *dsx*-GAL4>myrGFP (ere*: with the exception of *D. erecta*, where CsChrimson:tdTomato was used for labeling) across five species. san: *D. santomea*; tei: *D. teissieri;* ere: *D. erecta*. **(e)** Morphological subtypes of TN1 neurons and illustration of the core sine song circuit in *D. melanogaster* based on^23,31^. **(f)** Representative confocal stacks showing the anterior and posterior arbors of *dsx* neurons in the TN1 region across five species. **(g)** Relative volumes of the defined anterior arbors and posterior arbors highlighted one one side in panel **f**, *versus* all arbors in the TN1 region (AA/TN1 and PA/TN1) across five species. Data are presented as mean±SD with sample sizes shown in parentheses. We performed a one-way ANOVA with a Tukey-Kramer test for multiple comparisons (panels **c**,**g**) or a one-way Kruskal-Wallis test with a Bonferroni correction (panel **d**). Different letters in panels **d**,**g** denote significant differences (p < 0.05). Scale bars: 10 µm (panels **c**,**d**) and 50 µm (panel **f**).

Among species, song types and parameters are highly variable^14^. One notable lineage is *D. yakuba* and its sibling species *D. santomea*. They share a recent common ancestor with *D. melanogaster*, but have completely lost sine song and instead sing pulse and clack, two types of pulse-shaped songs that differ in waveform and social context^10,17,18^ (**Fig 1a**). In *D. melanogaster*, the production of sine song relies on the function of the sex determination gene *doublesex* (*dsx)* and *dsx*-expressing neurons^19–23^. Particularly, *dsx* is spliced into sex-specific isoforms that exert distinct developmental effects on neurons that serve key circuit functions for sine song generation.

Leveraging the lineage-specific loss of sine song in *D. yakuba*^24,25^ and the entry points provided by the characterized mechanisms of sine song generation in *D. melanogaster*^19–23^, we developed genetic tools that permit species comparisons at the circuit, cellular, developmental, and molecular levels to systematically investigate the mechanisms underlying behavioral evolution. We show that the loss of sine song results from evolutionary changes to the cellular makeup of motor patterning circuits, where *D. yakuba* lost the key neuronal subtype that acts as the central node of the sine-generating circuit. We further demonstrate the evolutionary role of *dsx*-dependent programmed cell death in altering the cellular makeup of song patterning neurons.

## Results

### The decision-making pathway for sine song likely exists in *D. yakuba* to drive pulse song

*Drosophila* males sing specific song types in a social context-dependent manner^10,16^. The absence of sine song in *D. yakuba* could, in theory, result from changes at any node within the circuitry implementing sensorimotor transformation from interpreting the social context to generating motor responses. When relatively close to a female and moving slowly in her direction, *D. melanogaster* males preferentially sing sine song, while *D. yakuba* males, who have lost the ability to sing sine, sing pulse song^10,17^. Because *D. melanogaster* sine and *D. yakuba* pulse are used in similar social contexts, we hypothesized that the same decision-making pathway that interprets the social context would underlie these two different song types, and therefore a hybrid male might sing both song types at the same time in the appropriate social context. In line with this hypothesis, hybrid males between *D. yakuba* and *D. teissieri*, its closest hybridable relative that still sings sine song, produced a *D. yakuba*-like pulse song with an inter-pulse interval that resembles *D. teissieri* sine song in waveform and carrier frequency (**Fig. 1b** and **Fig. S1a-c**). In this chimeric song, the two song types, *D. yakuba* pulse and *D. teissieri* sine, manifested in a time-locked manner while maintaining their distinct acoustic characteristics. This implies that *D. yakuba* pulse and *D. teissieri* sine are the activity of different motor programs elicited by the same decision-making pathway. Therefore, the decision-making circuit that triggers sine song is likely present in *D. yakuba* to drive pulse song, and the loss of sine song in *D. yakuba* possibly occurs in the downstream motor patterning circuits in the wing neuropil of the ventral nerve cord (VNC).

### *D. yakuba* has fewer sine-song-essential TN1 neurons

In the wing neuropil of the VNC, TN1 neurons, a cluster of ∼30 male-specific interneurons per hemisphere, are both sufficient and necessary for sine song generation in *D. melanogaster*^22,23,26^. Given the critical role of TN1 in sine song, we first investigated their presence in *D. yakuba*. TN1 neurons are one of the anatomically distinct *dsx*-expressing neuronal clusters^22,26^, allowing the quantification of TN1 cell number by probing *dsx* mRNA or protein. Both *in situ* hybridization chain reaction (HCR)^27,28^ using *dsx* mRNA probes and immunostaining (IHC) using Dsx antibodies^29^ confirmed the presence of a TN1 cluster in *D. yakuba*, but with ∼10 fewer neurons than in *D. melanogaster* (**Fig. 1c**). By CRISPR/Cas9-mediated gene targeting, we generated *dsx*-GAL4 alleles by introducing a GAL4 at the endogenous *dsx* locus to drive the expression of reporter genes in both species (**Fig. S2a**). The resultant alleles fully recapitulated previously reported *dsx* expression (**Fig. S2b,c**), the pattern of *dsx* mRNA and Dsx protein, as well as the species difference in TN1 number (**Fig. 1c,d**), demonstrating the fidelity of this knock-in approach to quantify TN1 neurons.

To determine if the fewer TN1 neurons in *D. yakuba* represents a loss event that phylogenetically correlates with the loss of sine song, we generated *dsx*-GAL4 alleles in three additional species using the same gene targeting design (**Fig. S2a**). The species that lost sine song, *D. yakuba* and *D. santomea*, have the fewest TN1 neurons. *D. teissieri*, who sings a sine song with a more irregular waveform^10^, has an intermediate number of TN1 neurons between *D. yakuba*/*D. santomea* and *D. melanogaster*. *D.erecta* sings a sine song more similar to *D. melanogaster*’s^10^ (**Fig. 1a**) and has a comparable number of TN1 neurons (**Fig. 1a,d**). Together, these data suggest a gradual loss of TN1 neurons across the phylogeny in a manner that correlates with the loss of sine song.

### *D. yakuba* has divergent TN1 neuroanatomy

TN1 is a heterogeneous population with five anatomical subtypes, among which the subtype TN1A plays an essential role in sine song generation^23,30–32^. TN1A neurons activate the downstream hg1 motoneuron and therefore the hg1 muscle required for sine song production^33^ (**Fig. 1e**). This functional connectivity relies on the posterior arbors of TN1A neurons^23^ that provide synaptic inputs into the hg1 motoneuron both directly and indirectly^31^. Reducing TN1A posterior arbors by ablating *dsx* activity during development impairs this functional connectivity and abolishes sine song^23^.

Because TN1A posterior arbors are essential for the functional connectivity that generates sine song^23^, we quantified the neurite volume of *dsx*+ neurons in the posterior base of the TN1 region across five species. In *D. melanogaster*, arbors in this region are primarily contributed by TN1A neurons^23^. *D. yakuba* and *D. santomea* possessed the sparsest posterior arbors (22.2% and 31.4% of the *D. melanogaster* level, respectively) (**Fig. 1f,g**). *D. teissieri* and *D. erecta* exhibited an intermediate level of posterior arbors (56.5% and 59.8%). In contrast, the volume of anterior arbors remains largely the same across species. Therefore, *D. yakuba* and *D. santomea* have experienced a major reduction of the TN1A-characteristic arbors required for the circuit connectivity generating sine song.

### TN1 neurons lack the ability to drive sine song in *D. yakuba*

Given the major reduction in the number of TN1 neurons and the TN1A-characteristic arbors in *D. yakuba*, we predict that its TN1 neurons lack the necessary anatomical foundation for the functional coupling with the hg1 motoneuron to drive sine song. To directly compare TN1 function across species, we referred to the genetic intersection that labels TN1 neurons in *D. melanogaster*^23^ and developed the corresponding genetic reagents (*dsx*-GAL4∩13H01-LexA) that specifically target the complete set of TN1 neurons in *D. yakuba* (**Fig. 2a**). The labeled TN1 neurons confirmed the striking species differences in TN1 neuroanatomy in the posterior region (**Fig. 2b,c**). Using the genetic reagents, we optogenetically activated TN1 neurons by expressing CsChrimson^34^ in *D. melanogaster* and *D. yakuba*. Consistent with the previous report^23^, TN1 activation in *D. melanogaster* males elicited unilateral wing extension accompanied with wing vibration that generates sine song, resembling the natural behavior. In *D. yakuba*, TN1 activation also elicited unilateral wing extension, but without wing vibration or song (**Fig. 2d** and **Supplementary Video 1**). Bilateral wing extensions, occasionally co-occurring with vibrations generating a buzzing sound, were observed mostly at higher activation levels. However, the combination of this bilateral wing posture and the sound does not match any natural behaviors and is likely artificial. Importantly, under the conditions that elicited unilateral wing extension, the natural wing posture of sine song, TN1 activation reliably induced sine song in *D. melanogaster* but never triggered any wing vibration nor song in *D. yakuba*, supporting a modular change of TN1 functionality in relation to the loss of sine song.

**Fig. 2.**
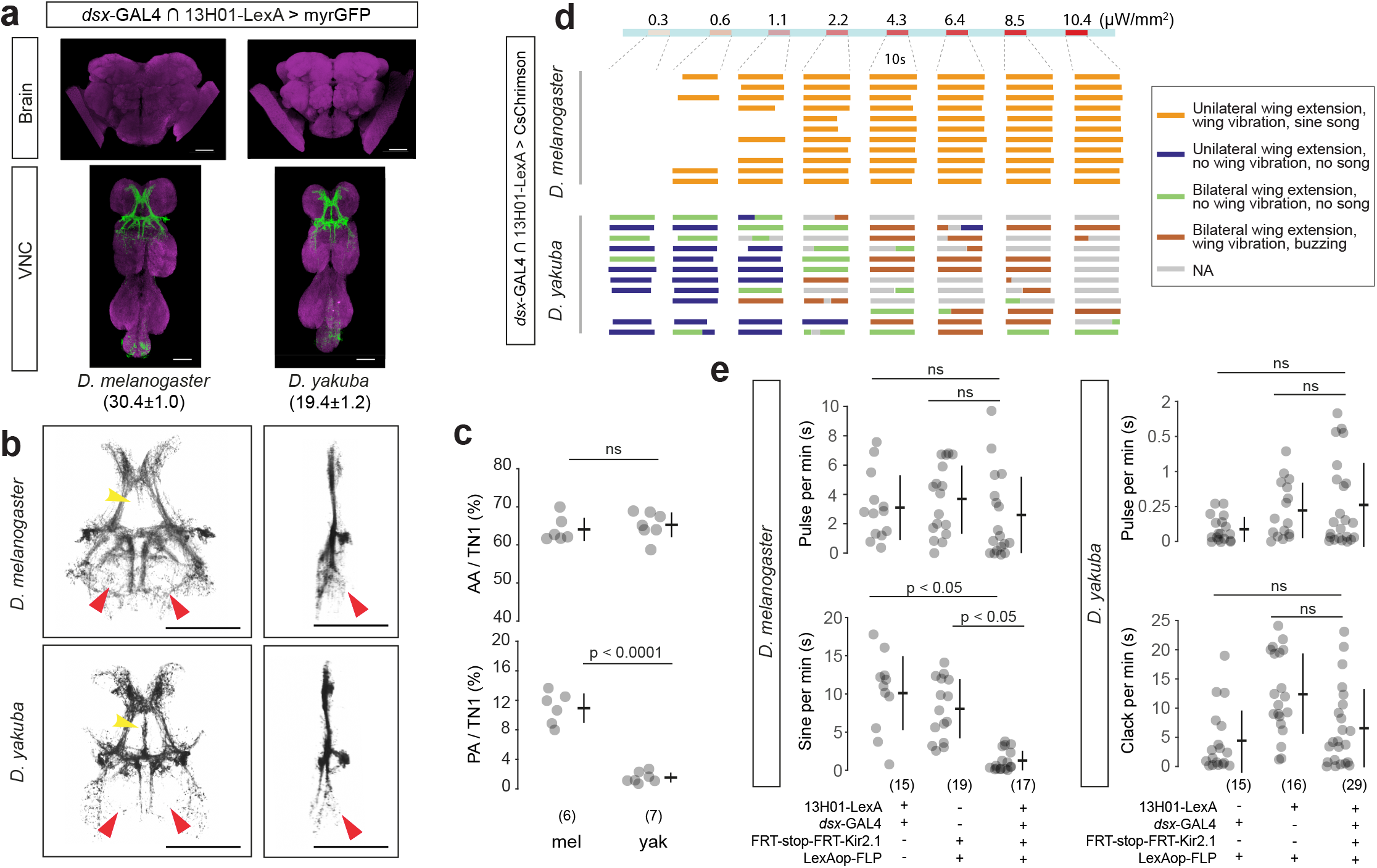
Functional comparison of TN1 neurons between *D. melanogaster* and *D. yakuba*. **(a)** The genetic intersection *dsx*-GAL4∩13H01-LexA specifically labels TN1 neurons in adult males similarly in *D. yakuba* as in *D. melanogaster*. Numbers of labeled TN1 neurons (per hemisphere) are noted in parentheses as mean±SD (n=10 for *D. melanogaster* and n=16 for *D. yakuba*). **(b)** Zoomed-in views (left: coronal; right: sagittal) of segmented TN1 arborizations in both species. Arrows denote two regions showing clear species differences: the posterior arbors (red) and the anterior medial arbors (yellow). **(c)** Relative volumes of the defined anterior arbors (AA) and posterior arbors (PA) *versus* all TN1 arbors in *D. melanogaster* (mel) and *D. yakuba* (yak). **(d)** Behavioral responses upon optogenetic activation of TN1 neurons at ramping intensities on a 10s ON and 20s OFF scheme (OFF period not displayed proportionally). Each row represents a male. Colors represent the defined behavioral categories. NA: not applicable to define. Also see **Supplementary Video 1**. **(e)** Song phenotypes of TN1 inhibition. Data are presented as mean±SD with sample sizes shown in parentheses. We performed a one-way ANOVA (panels **b**,**d**) and Mann-Whitney U test with a Bonferroni Correction (panel **e**). ns, not significant. Scale bars: 50 µm.

Across the phylogeny, the loss of sine song in *D. yakuba* occurred together with the gain of a second pulse-shaped song type, clack song^10,17,18^. Therefore, we tested whether the function of TN1 neurons has shifted in *D. yakuba* to generate clack or pulse. We expressed Kir2.1, an inwardly rectifying potassium channel^35^, in TN1 neurons for neuronal inhibition in *D. melanogaster* and *D. yakuba*. Similar to the previous report^23^, inhibiting TN1 neurons led to less sine song in *D. melanogaster*. We did not detect any significant effect on the amount of pulse or clack songs in *D. yakuba* (**Fig. 2e**). Thus, our data did not provide support for a major functional shift of TN1 neurons to mediate the gain of pulse-shaped song type in *D. yakuba*. We note that it is still possible that TN1 neurons are functional in *D. yakuba*, and as a general caution, using different effectors or more sensitive assays might reveal a phenotype.

### *D. yakuba* has lost the sine-song-essential TN1A neurons

The fewer TN1 neurons, the sparser TN1A-characteristic posterior arbors, and the inability of TN1 neurons to drive sine song in *D. yakuba* together imply a loss of TN1A neurons, the key TN1 subtype for sine song generation. Alternatively, TN1A neurons might still exist in *D. yakuba* with diverged morphology while retaining the overall molecular identity. Using a single-cell transcriptome approach, we compared the molecularly-defined subtypes of TN1 neurons between *D. yakuba* and *D. melanogaster*. We generated single-cell RNA sequencing (scRNAseq) data of genetically labeled and fluorescently sorted *dsx*+ neurons from adult male brains and VNCs in both species. By integrating the datasets from the two species and performing a clustering analysis^36^, we identified a putative TN1 cluster that expresses *TfAP-2*, a transcription factor that marks the 12A hemilineage origin of TN1 neurons^37^ (Fig. S3a). We also found such a *dsx* and *TfAP-2* co-expressing cluster using the published scRNAseq data of the adult male VNC in *D. melanogaster*^38^ (**Fig. S3b**). Employing HCR and genetic labeling, we verified that the co-expression of *TfAP-2* and *dsx* defines the complete set of TN1 neurons in male VNC in both species (**Fig. S3c-f**).

Next, we extracted the TN1 cluster from the full dataset for subclustering to resolve potential TN1 subtypes. Among the five identified molecular subclusters, four are present in both species, and one is absent in *D. yakuba* (**Fig. 3a**) – a pattern robust to varying clustering parameters (**Fig. S4a**). The gene expression patterns of selected key marker genes and neurotransmitters are conserved between species (**Fig. S4b**), and we also did not detect any species differences in the expression level of *dsx* (**Fig. S4c**). To confirm the *D. melanogaster*-specific TN1 subcluster and characterize its identity, we identified the molecular marker *CCKLR-17D3* (*17D3*) that is ubiquitously expressed in this subcluster, with sparse expression in the neighboring subcluster and no expression in the remaining subclusters (**Fig. 3b**). By joint-IHC-HCR, we detected an average of 8.1 *17D3*+ TN1 neurons in *D. melanogaster* and 1.3 in *D. yakuba* (**Fig. 3c**), confirming the loss of *17D3*+ TN1 neurons in *D. yakuba*. We hypothesized that the *17D3*+ TN1 neurons represent neurons of the anatomically defined subtype TN1A. Indeed, the entire *17D3+* TN1 population is labeled by a TN1A line in *D. melanogaster*^23^ (**Fig. 3c**). We further designed a genetic reagent that specifically labels *17D3*+ TN1 neurons, exploiting their co-expression of *dsx*, *TfAP-2*, and *17D3*. In *D. melanogaster*, the three-way intersection (*TfAP-2*-AD∩*dsx*-LexADBD∩*17D3*^GAL4^) labeled neurons resembling TN1A morphology featured by strong posterior arborizations (**Fig. 3d**). Inhibiting these neurons by expressing Kir2.1^35^ caused a reduction in sine song but not pulse song (**Fig. 3e**). Together, these data support a loss of sine song due to the loss of TN1A neurons in *D. yakuba*.

**Fig. 3.**
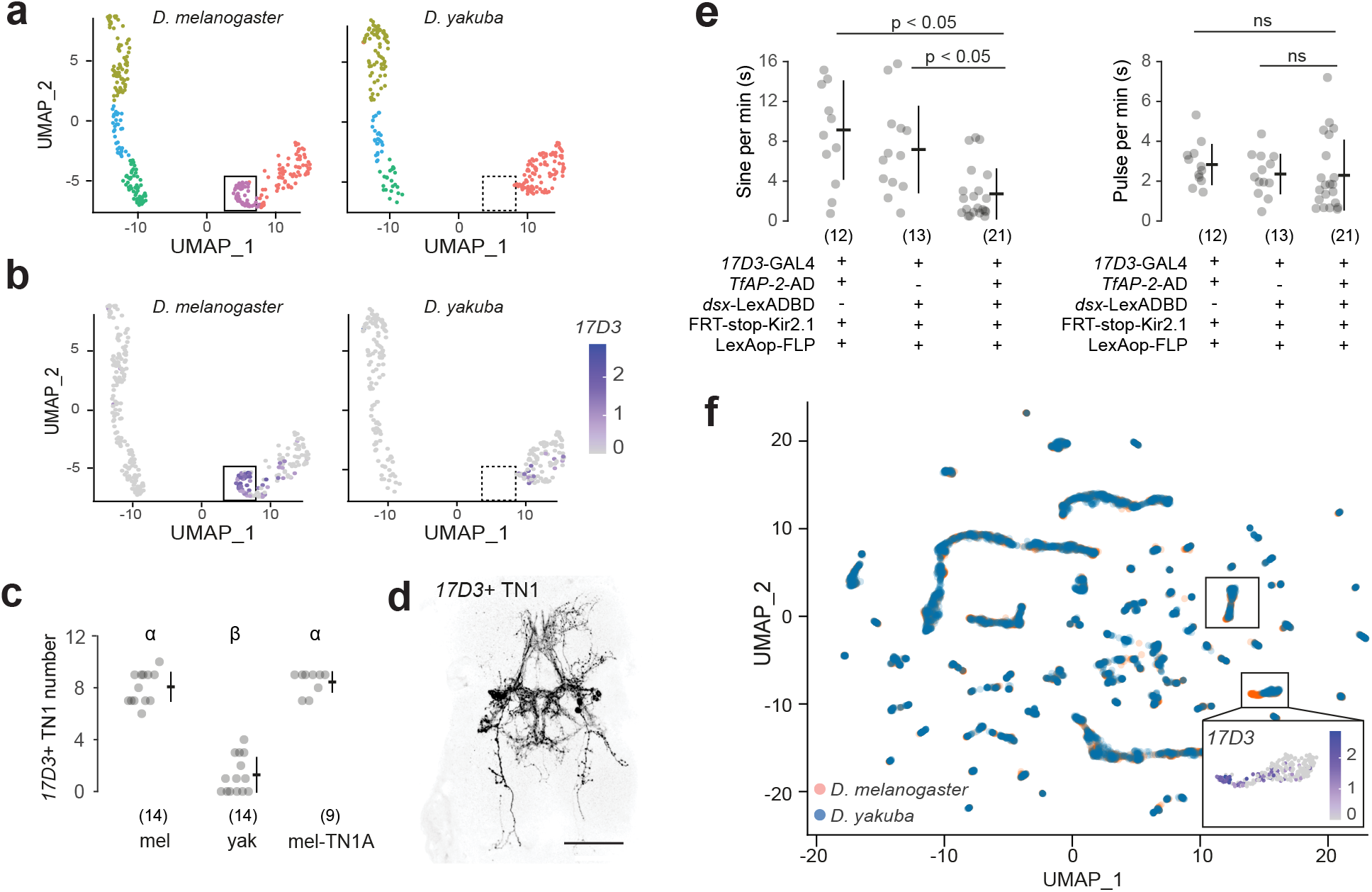
*D. yakuba* has lost a cell type that represents the sine-song-essential TN1A neurons. **(a)** UMAP representations of the scRNAseq data of TN1 neurons in *D. melanogaster* and *D.yakuba*. The box highlights the *D. melanogaster*-specific cluster. **(b)** Gene expression of *17D3*, a marker gene for the *D. melanogaster*-specific TN1 cluster, on the same UMAP representations as panel **a**. **(c)** Number of *17D3*+ TN1 neurons (per hemisphere), measured by joint-IHC-HCR, in *D. melanogaster* (mel), *D. yakuba* (yak) and a split-GAL4 line that labels TN1A neurons in *D. melanogaster* (mel-TN1A): VT017258-p65AD::Zp∩*dsx^Zp::Gdbd^*>myrGFP^23^. This split-GAL4 line labels 15.4±2.6 TN1 neurons. **(d)** Confocal stacks of *17D3*+ TN1 neurons labeled by the three-way genetic intersection *TfAP-2*-AD∩*dsx* LexADBD∩*17D3*^GAL4^>myrGFP. This intersection labels 5.9±1.1 neurons, with 4.8±0.8 neurons resembling TN1A and 1.1±0.9 neurons resembling TN1C based on the characteristic neural tracts. **(e)** Song phenotypes of inhibiting *17D3*+ TN1 neurons in *D. melanogaster*. **f**) UMAP representations of all *dsx*+ neurons in the brain and VNC for *D. melanogaster* and *D. yakuba*. *D. yakuba* cells are plotted on top of *D. melanogaster* cells to illustrate the *D. melanogaster*-specific cell population. Boxes highlight TN1 cells and the expression of *17D3* is shown on a zoomed-in view of the *D. melanogaster*-specific population. Data are presented as mean±SD with sample sizes shown in parentheses. We performed a one-way Kruskal-Wallis test (panel **c**) and Mann-Whitney U test with Bonferroni Corrections (panel **e**). Different letters in panel **c** denote significant differences (p < 0.05). Scale bars: 50 µm.

We further asked the extent to which closely related species differ in molecular cell types, and whether TN1 represents an exceptionally variable population. We performed a clustering analysis on the entire scRNAseq dataset of *dsx+* neurons, which include many populations playing various roles in sexual behaviors^39,40^. Among the 97 molecularly defined cell clusters of *dsx+* neurons, the TN1 cluster containing the *D. melanogaster*-specific *17D3*+ population is the only one with an obvious species difference in this clustering space **(Fig. 3f**). With the notion that more subtle differences in cell types might be present between the species, TN1A neurons appear as a prevailing distinction.

### *dsxM*-mediated TN1 cell death in *D. yakuba*

We next investigated the molecular and developmental mechanisms underlying the quantitative loss of TN1 neurons in *D. yakuba*. In *D. melanogaster*, *dsx* regulates TN1 development in a sex-specific manner depending on the splice isoforms^21,22^. The male transcript d*sxM* is required during development to increase the density of TN1A posterior arbors while the female transcript *dsxF* removes TN1 neurons by apoptosis^21–23^. Given the key role of *dsx* in TN1 development, we tested whether the species difference in TN1 cell number depends on the function of *dsxM*. In *D. melanogaster*, the loss of *dsx* function in *dsx* null males did not change the number of TN1 neurons labeled by *dsx*-GAL4. In contrast, the loss of *dsx* in *D. yakuba* resulted in ∼5 more TN1 neurons, suggesting that the *dsxM* function leads to fewer TN1 neurons in *D. yakuba* (**Fig. 4a**). Since *dsx* is not expressed in the relevant neural stem cell to affect cell proliferation or in the glial cells to have a cell-nonautonomous effect^21^, *dsxM* likely reduces TN1 neurons in *D. yakuba* by influencing neuronal survival in a cell-autonomous manner. As expected, blocking the apoptosis of *dsx+* neurons by expressing the effector gene P35^41^ increased the number of TN1 neurons to the level of the *dsx* null in *D. yakuba*. Expressing P35 in *dsx+* neurons in *dsx* null males did not further increase TN1 number, indicating that P35 and *dsx* act through the same developmental process (**Fig. 4a**). Consistent with the lack of an effect of *dsx* on TN1 number in *D. melanogaster*, we did not see an effect of P35 either. In comparison to the divergent function of *dsxM* between species in males, *dsxF* serves a conserved function in promoting TN1 apoptosis in females, where both the loss of *dsx* and expression of P35 led to a gain of TN1 neurons in *D. yakuba* (**Fig. S5a**) like in *D. melanogaster*^21^. Therefore, in *D. yakuba*, *dsxM* has acquired a *dsxF*-like function to reduce the number of TN1 neurons by promoting apoptosis. This functional change of *dsx* likely co-occurred with the loss of sine song, because the *dsx* null males in *D. santome*a also exhibited an increase of TN1 neurons (**Fig. 4a**).

**Fig. 4.**
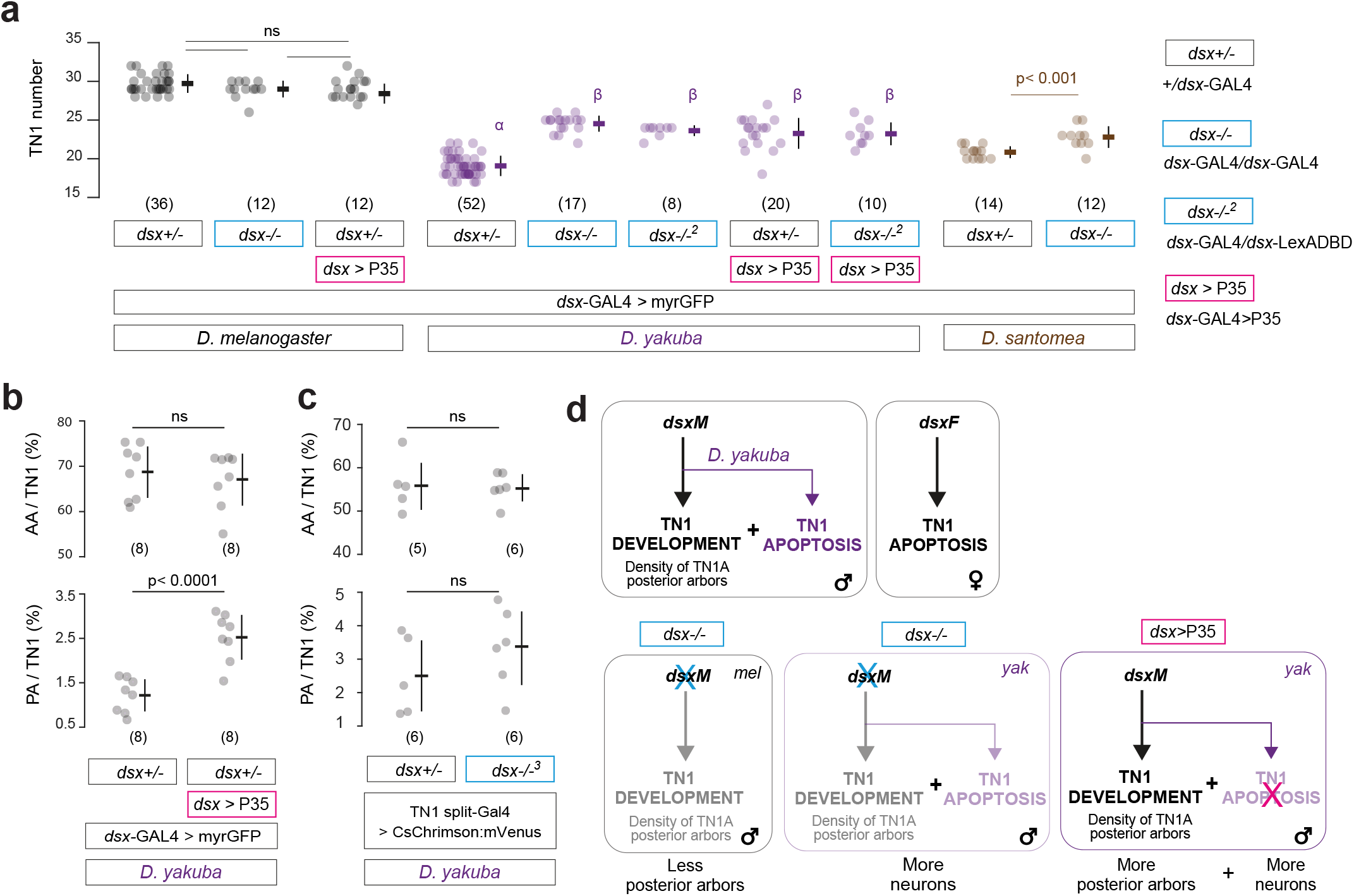
*dsxM* contributes to TN1 apoptosis in *D. yakuba*. **(a)** Effect on TN1 cell number (per hemisphere) in *dsx* null males (*dsx*-/-) and by blocking apoptosis in *dsx* neurons (*dsx*>P35) across species. Genotype abbreviations are specified on the right. **(b)** Effect on the relative volumes of TN1 anterior arbors (AA/TN1) and posterior arbors (PA/TN1) by blocking apoptosis in *dsx* neurons in *D. yakuba*. Volumes were measured based on the neurites in the TN1 region labeled by *dsx*-GAL4>myrGFP. Genotypes use the same abbreviations as panel **a**. **(c)** Effect on the relative volumes of TN1 anterior arbors (AA/TN1) and posterior arbors (PA/TN1) in *dsx* null males in *D. yakuba*. Volumes were measured based on the neurites labeled by TN1 split-GAL4 (*TfAP-2*-AD∩*dsx*-GAL4DBD)>CsChrimson:mVenus. *dsx*+/-: +/*dsx*-GAL4DBD; *dsx*-/-^3^: *dsx*-GAL4DBD/*dsx*-LexADBD. **(d)** Model for *dsx* function across species. The black pathway represents the conserved functions of *dsxM* in regulating TN1 development in males and *dsxF* in promoting TN1 apoptosis in females. The purple pathway represents the *D. yakuba*-specific function of *dsxM* in promoting TN1 apoptosis in males, a role that mimics the function of *dsxF* in females. Lower: summary of the resulting neuroanatomical outcomes in *dsx* null males and by blocking apoptosis based on the model. Data are presented as mean±SD with sample sizes shown in parentheses. We performed a one-way Kruskal-Wallis test (panel **a**) and Mann-Whitney U tests (panels **b** and **c**) with Bonferroni Corrections. Shared letters in panel **a** denote no significant difference (p > 0.05). ns, not significant.

In *D. yakuba*, blocking the apoptosis of *dsx* neurons resulted in about a twofold increase of the TN1 posterior arbors (**Fig. 4b**), potentially suggesting the resurrection of some TN1A neurons. In contrast, *dsx* null males, despite the resurrection of TN1 neurons, did not show an increase in the density of posterior arbors (**Fig. 4c**), consistent with a conserved role of *dsx* in the proper development of TN1A morphology^23^. Taken together, these findings support a dual-function of *dsxM* in *D. yakuba*: a species-specific one to reduce TN1 cell number, and a species-shared and potentially latent one that increases the posterior arbors of TN1A neurons if they exist (see model in **Fig. 4d**).

Lastly, we attempted to test if TN1 resurrection restores any song function in *D. yakuba*. The TN1 split-GAL4 reagent (*TfAP-2*-AD∩*dsx*-GAL4DBD), designed based on the molecular markers from the scRNAseq data, labels all TN1 neurons in the VNC and a cluster of neurons in the brain (**Fig. S3e**), allowing a feasible genetic scheme to specifically activate TN1 neurons in decapitated males with TN1 resurrection. However, this line could not drive P35 expression early or strong enough to sufficiently block TN1 apoptosis (**Fig. S5a,b**). Further, as expected, due to the lack of *dsx* activity (also see model in **Fig. 4d**), TN1 resurrection in *dsx* null flies did not restore sine song (**Fig. S5c**). Nevertheless, activating TN1 neurons in *dsx* null males was able to elicit unilateral wing extension with up-and-down wing flicking (appears as a slower motion of wing vibration) (**Fig. S5c** and **Supplementary Video 2**). This partial functional restoration leaves open the possibility of behavioral consequences due to the species difference of *dsxM* function in the developmental fate of TN1 neurons.

## Discussion

How evolution generates species diversity in complex motor behaviors is largely unknown. In this study, we leveraged the lineage-specific loss of sine song and integrated comparative approaches, from genes to circuits, to identify how evolution tinkers the biological system in behavioral evolution. First, we locate the neural substrates of evolution to the motor patterning circuits downstream of the decision-making pathway. Second, we link the loss of sine song with species differences in the cell number, neuroanatomy, and behavioral function of the song patterning neurons TN1. Third, we attribute the loss of sine song to the species difference in cell types, where *D. yakuba* has specifically lost neurons of the TN1A subtype essential for sine song generation. Fourth, we address the underlying developmental and molecular mechanisms, where the male isoform of the sex determination gene *dsx* in *D. yakuba* has a female isoform-like function to reduce TN1 number by promoting apoptosis.

Our results suggest a possible circuit principle underlying species diversification of motor repertories: shared decision-making pathways driving diverged motor outputs through the evolution of motor patterning circuits. The chimeric hybrid song, made of *D. yakuba* pulse and *D*. *teissieri* sine, indicates that the decision-making pathway for sine song is likely retained and repurposed in *D. yakuba* to drive pulse song. In parallel, the hybrid males sang a second song type resembling a chimera of *D. yakuba* clack and *D. teissieri* pulse (**Fig. S1**), echoing the previous finding that these two song types are driven by the same descending command neurons between species^10^. These data consistently imply selective pressures maintaining the decision-making pathways that interpret social contexts while diversifying their motor outputs, pointing to the contribution of motor patterning circuits, where TN1 neurons have experienced major changes in structure and function in relation to the loss of sine song. Also notable, the neuroanatomical differences of TN1 neurons are not specific to the comparison between *D. melanogaster* and *D. yakuba* but appear widespread among the five species we examined with divergent songs (**Fig. 1d,f,g**). Central pattern generators have been linked to species differences in rhythmic behaviors such as swimming in sea slugs and vocalization patterns in frogs and crickets^42–45^. Collectively, these findings support motor patterning circuits as common neural substrates mediating species diversification of behaviors.

The difference in TN1 subtypes among two closely-related species and its role in the loss of sine song contrasts with previous studies, which emphasize a conserved motor circuit organization^42,46^ and behavioral evolution through functional reconfiguration and synaptic rewiring of the underlying circuits^47–50^. Indeed, the loss of TN1A neurons as an evolutionary solution to remove sine song may appear counterintuitive given their recently identified role in pulse song in *D. melanogaster*. TN1A neurons are strongly connected to the core pulse-generating circuit and stay active during pulse singing; inhibiting them also affects a pulse song parameter^30,31^. However, compared to this relatively subtle role in pulse song, TN1A neurons serve as the central node of the core sine-generating circuit^31^ and are essential for sine song generation^23,30–32^. Recent connectome data further raises the possibility of functional heterogeneity within TN1A neurons, with subgroups that are more or less connected to the pulse-generating circuit^30^. Therefore, the functional indispensability and specialization of TN1A neurons in sine song generation make it a circuit node poised for evolutionary modification to yield major behavioral outcomes with limited pleiotropic effects. Reconciling with the general themes of highly interconnected nervous systems and widespread neural activity during specific behavioral modes^51^, a functionally modular organization of motor patterning circuits^52^, where some circuit nodes are tailored for specific motor patterns as seen in some complex motor systems, may provide the evolutionary flexibility needed to encode a diverse array of motor patterns by altering the cellular makeup of the underlying circuits.

In theory, the fewer TN1 neurons in *D. yakuba* may simply reflect a loss of sexual identity due to the loss of *dsx* gene expression, which could impair the development of TN1A posterior arbors necessary for the functional connectivity that generates sine song. Gene expression changes of master regulators in development, including sex determination genes such as *dsx*, are a major mechanism in morphological evolution^53,54^. While the loss of *dsx* gene expression remains a possible contributor, our results revealed that the species-specific function of *dsxM* in promoting TN1 apoptosis plays a substantial role. In both invertebrates and vertebrates, programmed cell death regulated by sex determination genes and hormonal pathways occurs extensively to orchestrate the ontogenetic changes and sexual dimorphism in circuits and behaviors during normal development^55,56^. For example, seasonal variation of hormonal controls and apoptosis are associated with the yearly remodeling of song circuits and song behaviors in adult male birds^57–59^. The striking evolutionary lability of *dsx* to redefine the developmental fate of neurons underscores the evolutionary significance of the interplay between sex determination genes and developmental programs to reconfigure the central circuits by fine-tuning the cellular makeup. Future work will determine if the functional divergence of *dsx* is mediated by genetic changes on this transcription factor itself or other genes such as its downstream targets.

Moving beyond the loss of TN1A neurons and the *dsxM*-dependent TN1 cell death in *D. yakuba*, the mechanisms contributing to the species difference of TN1 neurons are likely complex. For example, the anterior apex of TN1 neurons in *D. yakuba* exhibits dense medial arbors that are minimally identifiable in *D. melanogaster* (**Fig. 2b** and **Fig. S3e**). Additionally, blocking apoptosis in *D. yakuba* did not fully restore TN1 number to the level of *D. melanogaster* (**Fig. 4a**). Thus, other mechanisms, such as alterations of cell types via changes in temporal factors and terminal selector genes^60^, may act together to modify the song patterning neurons during evolution.

In conclusion, by developing genetic tools in non-model *Drosophila* species that enable species comparisons, our work provides insights into the circuit, cellular, developmental, and molecular mechanisms by which complex motor behaviors evolve and exemplifies how evolutionary changes are distributed at and translated across different biological levels during behavioral evolution.

## Materials and Methods

### Fly lines

Flies were maintained on cornmeal-agar-yeast medium (Fly Food B, Bloomington Recipe, Lab Express) at 23°C. The genotypes of all fly lines used in this study are listed in **Supplementary Table 1**.

### Generating CRISPR/Cas9-mediated knock-in alleles and other transgenic lines

*dsx-*GAL4 (*D. melanogaster, D. yakuba, D. santomea, D. teissieri, D. erecta*): The *dsx*-GAL4 knock-in alleles were generated by replacing the majority of the first coding exon of *dsx* in-frame with GAL4 separated by the sequence of a T2A self-cleaving peptide. The donor plasmids were constructed by concatenating a left homology arm, T2A, GAL4 (amplified from pBPGUw, addgene #17575^61^), a Mhc-DsRed marker^62^, a right homology arm, and a 1.8 kb backbone using Gibson Assembly (NEB). T2A was initially introduced using a gBlock (IDT) that covers the left homology arm and T2A. Once the first donor plasmid was made, the T2A-GAL4 fragment can be amplified by PCR to assemble the rest donor plasmids. *dsx-*LexADBD (*D. melanogaster, D. yakuba*)*, dsx-*GAL4DBD (*D. yakuba*): The *dsx*-LexADBD *and dsx-*GAL4DBD knock-in alleles were generated using the same design as the *dsx-GAL4* alleles. The GAL4DBD fragment was amplified from the pBPZpGAL4DBDUw plasmid (addgene #26233). The LexADBD fragment was synthesized using a gBlock (IDT) based on the sequence of the plasmid pSW1324 (provided by Barry Dickson). All the resultant *dsx* alleles have the majority of the coding sequence deleted (**Fig. S2a**), therefore should function as null alleles.

*TfAP-2-*AD (*D. melanogaster and D. yakuba*): The basic design of the *TfAP-2* knock-in alleles was similar to those of the *dsx* knock-ins. The T2A-AD cassette was inserted into the first shared exon by all isoforms (the third exon) of *TfAP-2*, while generating an 87-bp deletion of the coding sequence at the insertion site.

For all CRISPR/Cas9-mediated knock-ins, a pair of guide RNAs (gRNAs) were cloned into the pCFD4 vector. The gRNAs were designed using DRSC CRISPR finder (https://www.flyrnai.org/crispr/), prioritizing those with high efficiency scores (>7) and starting with a guanine. A cocktail containing the donor plasmid (200 ng/ul), pCFD4 plasmid (200 ng/ul), and *in vitro* transcribed Cas9-nanos 3’UTR mRNA (200 ng/ul) was injected into fly embryos by Rainbow Transgenic Flies. The knock-in flies were screened using the Mhc-DsRed marker that drives DsRed expression in muscles. Detailed information on gRNA sequences and primer sequences was provided in **Supplementary Table 2**.

To generate UAS-P35 in *D. yakuba*, the coding sequence of P35 was amplified from the genomic DNA of UAS-P35^BH1^ (BDSC #5072) flies and inserted into the XbaI and XhoI sites of pJFRC81-10XUAS-IVS-Syn21-GFP-p10 (addgene #36432) via Gibson assembly (NEB). The resultant plasmid was injected into the *D. yakuba* attP line 2180^63^. Efficacy of the transgene P35 on apoptosis was confirmed in *D. yakuba* females, in which *dsx*-GAL4>UAS-P35 resurrected 4.6±3.5 TN1 neurons (**Fig. S5a**). The rest of the transgenic lines were generated by injecting published vectors. The vectors were introduced into the genomes using attB/P φc31 integrase system or piggyBac transgenesis^64^. All injections were performed at Rainbow Transgenic Flies using a standard protocol. Detailed information on the plasmids and the injected strains was provided in **Supplementary Table 3**.

### IHC, HCR, and joint-IHC-HCR

For IHC, brains and VNCs from 3-7 day old adults were dissected in cold 1X PBS for 50 min and then transferred to 4% paraformaldehyde (buffered in PBS) and fixed for 35 min at room temperature (RT). Samples were then rinsed 3 times in PBTX (1X PBS with 1% Triton X-100), and blocked in 5% NGS solution (buffered in PBTX) for 1.5 hours. We incubated the samples with primary antibodies diluted in 5% NGS solution (buffered in PBTX) overnight at 4°C, washed 3×30 min in PBTX at RT, and then incubated with secondary antibodies diluted in 5% NGS (buffered in PBTX) overnight at 4°C. The next day, after being washed 3×30 min in PBTX at RT, samples were mounted on poly-L-lysine coated coverslips with ProLongTM Gold antifade reagent (Fisher Scientific; Cat.#: P36931). Imaging was performed on a Leica DMi8 microscope with a TCS SP8 Confocal System or a Zeiss LSM710 confocal microscope system at 40x with optical sections at 0.4-0.8 µm intervals. Data for direct comparison were collected with the same setting on the same platform.

HCR^27^ was conducted following the “HCR RNA-FISH: Generic Sample in Solution’’ protocol provided by Molecular Instruments. Brains and VNCs were dissected, fixed, and blocked under the same conditions mentioned above. We then pre-hybridized samples in pre-warmed Probe Hybridization Buffer (Molecular Instruments) for 30 min at 37°C. Probe solution (16 nM, in pre-warmed Probe Hybridization Buffer) was applied for overnight incubation (∼16 hours) at 37°C. After hybridization, samples were washed with pre-warmed Probe Wash Buffer (Molecular Instruments) 4×15 min, and then with 5X SSCT (Saline-Sodium Citrate Solution with 0.1% Tween-20) 3×5 min. During these washes, DNA hairpin amplifiers (30 pmol) were digested at 95°C in a pre-heated thermal block for 1.5 min. Activated hairpins were then photo-protected and left to cool for 30 min while Amplification Buffer (Molecular Instruments) was applied to the samples for pre-amplification during the same time period. A hairpin solution was created by mixing the activated amplifiers with Amplification Buffer and applied to the samples for overnight incubation (∼16 hours) at RT. The following day, we washed the samples with 5X SSCT via a 2×5 min, 2×30 min, and 1×5 min series. Samples were mounted similarly as in IHC (Fisher Scientific; Cat.#: P36931).

Joint-IHC-HCR used an adapted version of a previously published protocol^28^. Following standard dissection and fixation, samples were blocked using Antibody Buffer (Molecular Instruments) for 4 hours at 4°C. We diluted the appropriate primary antibodies in Antibody Buffer and incubated the samples overnight (∼16 hours) at 4°C. The next day, following a 4×30 min PBTX wash series, samples were incubated for 3 hours in secondary antibodies diluted in Antibody Buffer at RT. Another wash series consisting of a 5×5 min PBTx wash and a 1×5 min 5X SSCT wash was completed before post-IHC fixation in 4% paraformaldehyde solution for 10 min. Samples were then rinsed with PBTX once and 5X SSCT twice. Excess probes were washed away using pre-warmed Probe Wash Buffer (Molecular Instruments, Inc.) 4×5 min followed by a 5 min 5X SSCT wash. From this point on, the HCR portion of joint-IHC-HCR was followed exactly as an independent HCR had been described. Detailed information on antibodies, mRNA probes, and hairpins were provided in **Supplementary Table 4**.

### Cell number and neurites quantification

Cell numbers were quantified from confocal images using the Fiji/ImageJ plugin “Cell Counter”. Neurite volumes were quantified using VVD Viewer (https://github.com/takashi310/VVD_Viewer). Starting with the dorsal view of the ventral nerve cord, putative TN1 arborizations were manually segmented with the function of “Paint Brush” on “Analyze” panel, and the numbers of pixels were counted with the function of “Analysis”. Then, the anterior apex and posterior base of segmented TN1 arborizations were selected and analyzed by the same procedure. For the posterior arbors, we only considered the neurites extending from the medial to the distal sides of the VNC and excluded the medial arbors, which were heavily contributed by non-TN1 *dsx+* neurons in *dsx*-GAL4>myrGFP flies. For the genetic reagents that label TN1 neurons specifically, we defined the anterior and posterior arbors in the same way to keep the comparability among data. To control for differences across species in VNC volume and labeling intensity, we calculated the volumes of the anterior and posterior arbors relative to the volume of the TN1 neurons in each sample.

### Behavioral experiments

Unmated males were collected and single-housed for 4-6 days at 23°C and 50% humidity with a 12-hr light/dark cycle. For non-optogenetic recordings, each male was loaded into a 5 mm diameter chamber with an aspirator to pair with a 1 day old, group-housed unmated female, and recorded for 30 min within the first 2-3 hours of the day cycle. Audio (5000 Hz) and video (10 Hz) recordings were simultaneously recorded on *Song Torrent*, a custom 96-channel recording apparatus^65^. For this setup, each chamber sits on a single microphone, and two cameras (FLIR BFS-U3-200S6M-C, Edmud optics #11-521) with 50mm lens (Edmud optics #63-248) capture the entire field.

For optogenetic experiments, unmated males were single-housed and reared on all-trans-retinal food (0.2 mM) for 4-6 days. Each male was loaded into a 5 mm chamber without a female. CsChrimson was stimulated with programmable red LEDs (635 nm), mounted on an aluminum rack facing down towards the chambers. Light stimulation was applied on a 10s on, 20s off cycle with 8 levels ramping from 0.3 to 10.4 μW/mm^2^ t. During the recordings, chambers were tonically illuminated with blue LEDs (STN-BBLU-A3A-10B5M-12V, Super Bright LEDs). Infrared LEDs (850 nm) were provided as a light source for video recording, where a lens was attached with an 800 nm long pass filter (Edmund optics #66070) to block optogenetic illumination. For optogenetic experiments using decapitated flies, males were anesthetized with ice, decapitated using dissecting scissors, and left to recover in the chambers for 15-20 min. Flies that failed to recover or keep a standing position were excluded from further experiments. The rest of the recording procedure followed the same protocol as described above. For both activation and deactivation experiments using *D. yakuba* flies, IHC was performed to confirm that flies express CsChrimson (via the fused mVenus) or Kir2.1 (via the fused GFP) as expected.

### Courtship video and audio analyses

Courtship songs of *D. melanogaster* were predicted and analyzed as previously reported^23,66^. Courtship songs of *D. yakuba* were predicted using SongExplorer, a deep learning-based interface for the discovery and segmentation of acoustic signals^67^. We manually annotated ground-truth data for three categories of events: “pulse”, “clack”, and “nosong” (“nosong” includes both ambient noise and non-song sound). A classifier was trained on the ground-truth data and applied for song prediction. We identified and sampled false predictions for correction using the SongExplorer functions “fix false positives” and “fix negative positives”, and then re-trained the classifier with ground-truth including these corrected values. This process was iterated until expected performance was achieved. The final ground-truth dataset includes 723 events for “pulse”, 1277 for “clack”, and 1488 for “nosong”. The prediction accuracy was 94.5% for “pulse”, 95.2% for “clack”, and 93.8% for “nosong” based on the confusion matrix. Ethograms were generated based on the predicted probabilities for each category using a threshold derived from the precision-recall curve and a precision-recall ratio of 1. Manual inspection of ethograms suggested that false positives were mostly contributed by events with low probability values that transiently rise above the threshold. Therefore, heuristics were applied to exclude predictions that last less than 0.02 millisecond. The predicted events were re-centered by the signal peak with a 0.15 millisecond window on each side and analyzed as previously reported^10,66^.

Video recordings of optogenetic activation were manually annotated using Tempo (https://github.com/JaneliaSciComp/tempo). Six categories of behaviors were defined and manually annotated: (1) unilateral wing extension, wing vibration, sine song; (2) unilateral wing extension, no wing vibration, no song; (3) bilateral wing extension, no wing vibration, no song; (4) bilateral wing extension, wing vibration, buzzing; (5) NA (cannot be scored because flies fell over); (6) unilateral wing extension, wing flicking (up and down wing movement in a slower motion of the typical wing vibration that generates song).

### scRNAseq libraries

For both *D. melanogaster* and *D. yakuba*, *dsx*-GAL4 females were crossed with males carrying UAS-nls::tdTomato to generate offspring expressing nls::tdTomato in *dsx+* neurons. Males were collected within 12 hours of eclosion and housed in groups of 20 at 23°C and 50% humidity with a 12-hr light/dark cycle. The brain and VNC of 4-8 day old males were dissected within 50 min in cold Schneider’s medium to prepare ∼100 flies per sample (900 *dsx+* neurons per fly x 100 flies for ∼90,000 *dsx*+ neurons per sample). Cells were dissociated following a modified version of a previously described protocol^68^. In short, the brains and VNCs were transferred into 5 ml Liberase DH (1 mg/ml, Supply Solutions), dissolved in Schneider’s medium, and digested on a rotator at RT for 45 min. When the digestion was finished, the sample was spun down and the dispase was removed with a pipette before washing with Schnieder’s medium three times. Next, 500 ul of 2% BSA in 1X PBS was added to the sample. The tissue was broken apart by pipetting up and down 100 times using a 200 ul pipette tip, pre-coated with 100% FBS to prevent cells from sticking. To further break apart the cells, a 3 cc/ml needle, pre-coated with 100% FBS, was used to pipette the solution an additional 20 times. Finally, the solution was transferred to a flow cytometry tube. Following dissociation, nls::tdTomato+ cells were sorted out by a fluorescence-activated cell sorting Aria Cell Sorter, using a 100 um nozzle and 1.25 drop precision (prior testing was performed to compare the fluorescence signals of sorting using myrGFP *versus* nls::tdTomato, and the latter shows a much cleaner separation of nls::tdTomato+ population from the rest in both species).

23.5% (*D. melanogaster*) and 27.1% (*D. yakuba*) of the initial ∼90,000 *dsx*+ cells were collected by cell sorting in each sample. To verify the yield of nls::tdTomato+ cells measured by sorting, the number of nls::tdTomato+ cells was counted on a disposable hemocytometer using a Leica M165 fluorescent scope. Upon confirming, the cells were processed at the Center for Applied Genomics at the Children’s Hospital of Philadelphia’s Research Institute where all library preparations and sequencing were performed. scRNA-seq libraries were generated using the 10x Genomics Single Cell 3’ Library and Gel Bead Kit following the manufacturer’s recommendation protocol^69^ and the libraries were sequenced on an Illumina NovaSeq 6000. The amount of time between the start of cell dissociation and the start of cell sorting was ∼70 min, sorting the cells lasted ∼30 min, and the time between the end of cell sorting and the start of the 10x Genomics protocol was ∼30 min.

### scRNAseq data analyses

The 10x Genomics CellRanger software v7.1.0 “count” command was used to map sequencing reads to the *Drosophila melanogaster* (Assembly GCA_000001215.4) and *Drosophila yakuba* (Assembly GCA_016746365.2) genomes, respectively. CellRanger filters cells based on the inflection point of the barcode-rank plot. After CellRanger’s filtering, 5132 *D. melanogaster* and 5834 *D. yakuba* cells were included in the filtered expression matrix. 92.9% of the *D. melanogaster* reads mapped to the genome and the sample had a total of 207,369,182 reads (average of 39,986 reads per cell), 2227 median genes per cell, and a sequencing saturation of 67.7%. 90.3% of the *D. yakuba* reads mapped to the genome and the sample had a total of 178,751,749 reads (average of 30,405 reads per cell), 2141 genes per cell, and a sequencing saturation of 61.1%.

Seurat v5.0.1 was used to analyze expression matrices^36^ and the default parameters on all Seurat functions were used unless otherwise stated. Matrices were filtered to retain only high-quality cells by removing cells with greater than 15% mitochondrial transcripts, fewer than 200 genes detected, or expressing more than 4000 genes. To identify orthologous genes between species, a reciprocal best hits BLAST was performed with default settings. 11,739 orthologous genes were identified between *D. melanogaster* and *D. yakuba* and only these genes were included in the subsequent analyses. The *D. melanogaster* and *D. yakuba* datasets were integrated using the Seurat functions “SelectIntegrationFeatures”, “FindIntegrationAnchors”, and “IntegrateData”. The functions “NormalizeData” and “ScaleData” were used to normalize and scale the data after integration. A principal components analysis (PCA) was performed and the number of statistically significant PCs were calculated using a Jackstraw analysis. The functions “runUMAP”, “FindNeighbors”, and “FindClusters” were used to reduce the dimensionality of the dataset and perform clustering. Clusters where the majority of cells lacked detectable *dsx* expression were removed, resulting in a total of 4691 *D. melanogaster* and 5219 *D. yakuba* cells in the final dataset. This represents 5.2-5.8x sequenced cells per neuron of adult male *dsx+* neurons. By estimation, 21.5-22.1% of the dissected *dsx+* neurons were recovered after cell sorting, and 5.2-5.8% of the *dsx+* neurons were recovered after sequencing.

To identify TN1 neurons from this larger dataset, cells that express any of the VNC markers (*Antp*, *Ubx*, *abd-A*, and *Abd*-*B*) were extracted first. We performed a cluster analysis on this VNC-only dataset and identified two putative TN1 clusters that expressed *TfAP-2*. We confirmed the intersection of *dsx* and *TfAP-2* labels TN1 by extracting all cells expressing *dsx* from the publicly available dataset from the single-cell transcriptomic VNC atlas^38^ and performing a cluster analysis to find that cells expressing *TfAP-2* also cluster together in this dataset. Finally, we performed a joint-IHC-HCR and genetic intersection to confirm that the combination of *dsx* and *TfAP-2* expression labels the entire TN1 population.

Upon confirmation, we extracted the *dsx* and *TfAP-2* co-expressing clusters from the larger dataset and performed another cluster analysis to better investigate variation within TN1 neurons. A wide range of PCs and clustering resolutions were compared to understand the robustness of the clustering. Conserved marker genes, genes with significantly higher expression in one cluster compared to all other clusters in both *D. melanogaster* and *D. yakuba* neurons, were identified for all clusters using the Seurat “FindConservedMarkers” function.

### Statistical Analysis

Data analysis was performed in R (v4.3.2) with the following packages: Seurat (v5.0.1), Matrix (v1.6.4), Seurat (v5.0.1), SeuratObject (v5.0.1), and SeuratDisk (v0.0.0.9021). Scripts are available by request. Before performing parametric tests (t-tests or ANOVAs), we checked whether the data were normally distributed using Shapiro-Wilk tests. If the data were not normally distributed, we performed nonparametric Mann-Whitney U or Kruskal-Wallis tests. Where relevant, we used Tukey-Kramer or Bonferroni corrections to adjust p-values for multiple comparisons.

## Acknowledgements

We thank Hongjie Li, Lihua Wang, Jean Rosario, and Junhyong Kim for sharing protocols and helpful discussions on scRNAseq. We thank David Stern and Hiroshi Shiozaki for sharing genetic reagents, Gerry Rubin and Heather Dionnefor for sharing plasmids and generating genetic reagents, and Yufeng Pan for sharing the DsxM antibody. We thank Troy Shirangi, Josh Lillvis, Rory Coleman, and Dawn Chen for comments on the draft. We thank Steven Sawtelle for helping with setting up the *Song Torrent* system and Benjamin Arthur for discussing audio analysis. This project was supported by Searle Scholarship and NIH grant R35GM142678 to Y.D.

## Author contributions

Y.D., D.Y., and J.T.W.conceived the study and wrote the paper. D.Y. and I.P.J. collected the confocal imaging and behavioral data. J.T.W. generated the scRNAseq data. D.Y. and J.T.W. performed data analyses. Y.D. and D.Y. generated genetic reagents.

## Figure Legends

**Fig. S1. Inferring the homology of song components between *D. yakuba*, *D. teissieri*, and their male hybrids.**

**(a)** Representative song traces showing the two song types in *D. yakuba* (yPulse and yClack), *D. teissieri* (tSine and tPulse), and the two chimeric song types in the F1 hybrid males. The first chimeric song type appears to be a pulse song (hPulse) with an inter-pulse interval (hIPI) resembling tSine. The second hybrid chimeric song type appears to be clack song (hClack) followed by a tail resembling a truncated train of tPulse. **(b)** Randomly selected examples of song components defined in panel **a**. Notably, hIPI shares the most similar waveforms with tSine. **(c)** Histogram of carrier frequencies for the defined song components. Number of total events and fly individuals (in parentheses) are noted on the right. Again, hIPI shares the most similar carrier frequencies with tSine. Based on the comparison of waveforms and carrier frequencies, we inferred the following homologous relationships among song components: yPulse is homologous to hPulse; tSine to hIPI, yClack to hClack, tPulse to hCT.

**Fig. S2. Generation of *dsx*-GAL4 knock-in transgenes.**

**(a)** CRISPR/Cas9-mediated targeting strategy of generating *dsx*-GAL4 alleles in *D. melanogaster*, *D. yakuba*, *D. santomea*, *D. teissieri*, and *D. erecta*. **(b)** Representative confocal images of *dsx*-GAL4>myrGFP in each species. **(c)** Confocal images of *D. melanogaster* and *D. yakuba* highlighting the cell bodies of previously defined *dsx*+ clusters^39,40^ in different colors. All *dsx*+ clusters can be unambiguously defined in both species based on the location of cell bodies and arborization patterns.

**Fig. S3. Identification of a conserved molecular marker for TN1 neurons in *D. melanogaster* and *D. yakuba*.**

**(a)** UMAP representations of molecular cell types in *dsx*+ neurons that express VNC markers (left) and the corresponding gene expression of *TfAP-2* (right). The circled cluster represents the putative TN1 neurons. **(b)** UMAP representations of molecular cell types in *dsx*-expressing neurons extracted from the published VNC scRNAseq dataset^38^ (left) and the corresponding gene expression of *TfAP-2* (right) in *D. melanogaster*. The circled cluster represents the putative TN1 neurons. **(c)** Confocal stacks showing the co-expression of *dsx* and *TfAP-2* in TN1 neurons (cell bodied circled) and the absence of *TfAP-2* expression in non-TN1 *dsx* neurons (arrowhead) in the regions of accessory mesothoracic neuropil and mesothoracic neuromere. **(d)** Co-expression of *dsx* and *TfAP-2* in the TN1 region is conserved between *D. melanogaster* and *D. yakuba*. **(e)** In VNCs of both species, the genetic intersection *TfAP-2*-AD∩*dsx*-GAL4DBD specifically labels TN1 neurons and reveals similar species differences in the posterior arbors (red arrows) and the anterior medial arbors (yellow arrows) as shown in Fig. 2b. This intersection also labels pC2l neurons in the brain. **(f)** The genetic intersection *TfAP-2*-AD∩*dsx*-GAL4DBD labels the complete set of TN1 neurons in both *D. melanogaster* (mel) and *D. yakuba* (yak). Data are presented as mean±SD with sample sizes shown in parentheses. We performed a one-way ANOVA. Scale bars: 50 uM.

**Fig. S4. Species comparison of TN1 neurons in molecular subtypes and expression of key genes based on scRNAseq data.**

**(a)** UMAP representations highlighting the *D. melanogaster*-specific cluster across a range of principal components and Seurat “FindClusters” function resolutions in *D. melanogaster* and *D. yakuba*. **(b)** UMAP representations split by species showing the expression levels of *elav* (neuronal marker), *dsx*, *fru* (sex determination genes), *TfAP-2*, *Antp* (TN1 marker genes), *ChAT*, *VAChT* (cholinergic neuron markers), *Gad1*, *VGAT* (GABAergic neuron markers), and *VGlut* (glutamatergic neuron markers). **(c)** Violin plots showing the expression levels of *dsx* in TN1 neurons between *D. melanogaster* and *D. yakuba*. We performed a Mann-Whitney U test, adjusted p-value = 1. Each dot represents a single-cell transcriptome of a TN1 neuron.

**Fig. S5. Additional data for TN1 resurrection in *D. yakuba*, related to Fig. 4**.

**(a)** P35 expression driven by the TN1-specific genetic intersection (TN1 split-GAL4>P35) in females had little effect in resurrecting TN1 neurons compared with *dsx* null (*dsx*-/-) and P35 expression driven by *dsx*-GAL4(*dsx*>P35). Genotype abbreviations are specified on the right. **(b)** Similarly, TN1 split-GAL4>P35 in males did not fully recapitulate the effects of *dsx* null (*dsx*-/-^3^) or *dsx*-GAL4>P35 (see Fig. 4a). This suggests that the TN1 split-GAL4 may not drive P35 sufficiently high or early enough to fully block apoptosis as *dsx*-GAL4 did. *dsx*+/-: +/*dsx*-GAL4DBD; *dsx*-/-^3^: *dsx*-GAL4DBD/*dsx*-LexADBD. **(c)** Behavioral responses of TN1 activation in decapitated males for the indicated genotypes. Unilateral wing extension coupled with wing flicking, albeit no song, was observed in *dsx* null males. Also see **Supplementary Video 2**. This result is consistent with its gain of TN1 neurons and the lack of *dsx* expression. The lack of a phenotype in TN1 split-GAL4>P35 flies is consistent with the insufficient resurrection of TN1 neurons. Data are presented as mean±SD with sample sizes shown in parentheses. We performed a Mann-Whitney U test with a Bonferroni Correction. Shared letters in panels **a**,**b** denote no significant difference (p > 0.05).

## Supplementary Information

**Supplementary Video 1.** Behavioral phenotypes of TN1 activation in *D. melanogaster* and *D. yakuba*, related to Fig. 2e. Under the conditions that elicited unilateral wing extension, TN1 activation triggered robust wing vibration coupled with sine song in *D. melanogaster*, but no wing vibration or song in *D. yakuba*. Red dots denote the activation periods.

**Supplementary Video 2.** Behavioral phenotype of TN1 activation in *dsx* null males in *D. yakuba*, related to **Fig. S5c**. Under the conditions that elicited unilateral wing extension, TN1 activation in *dsx* null males triggered wing flickings that appear as a slower motion of wing vibration. Red dots denote the activation periods. Males were decapitated to exclude the effect of brain neurons labeled.

**Supplementary Table 1.** *Drosophila* genotypes used in this study.

**Supplementary Table 2.** Primers to generate constructs for CRISPR/Cas9-mediated gene targeting.

**Supplementary Table 3.** Sources of *Drosophila* lines in this study.

**Supplementary Table 4.** Sources of antibodies, probes, and hairpins used in IHC and HCR.

