## Supplementary Figures and Tables for "Changes in the cellular makeup of motor patterning circuits drive courtship song evolution in *Drosophila*"

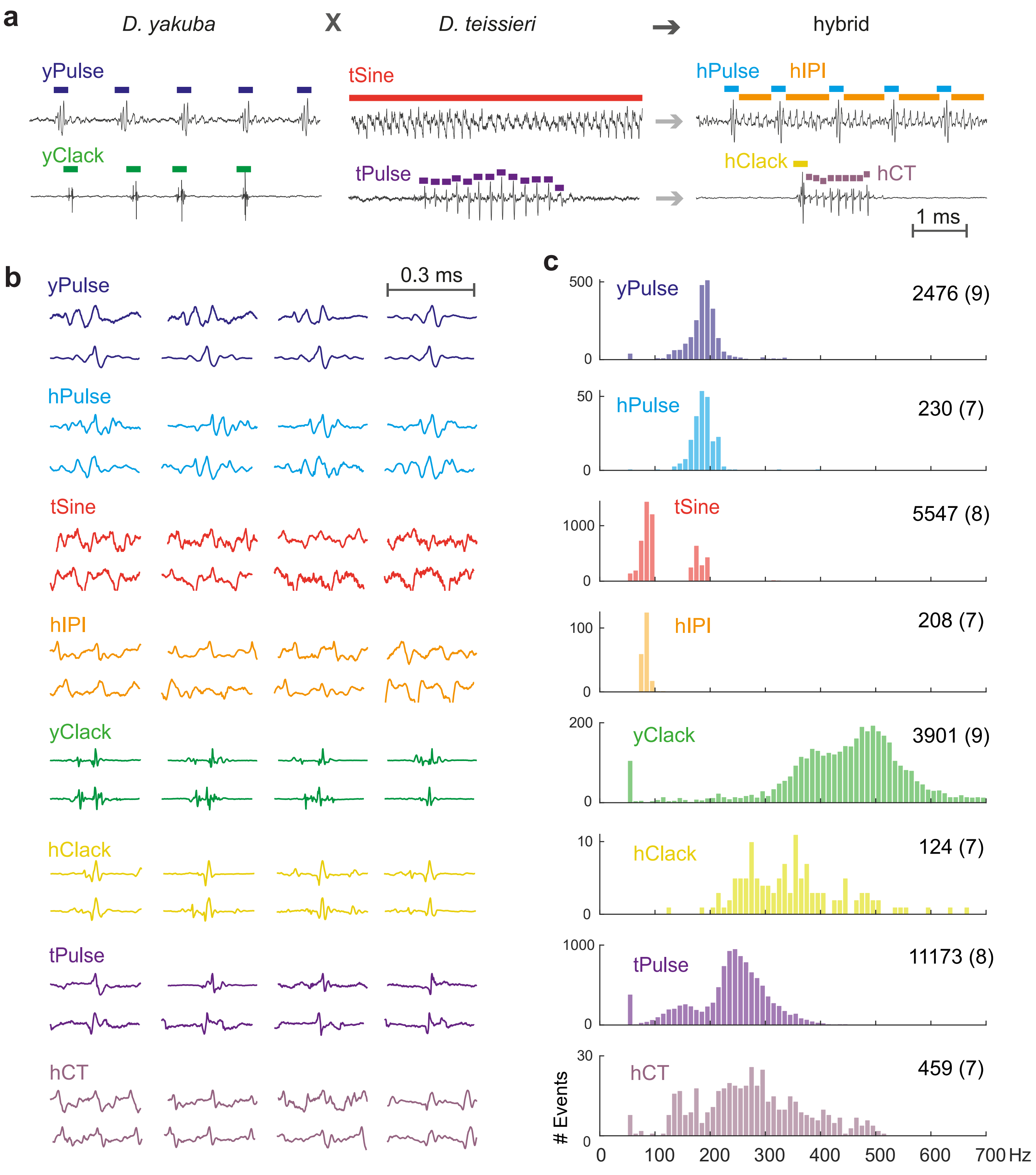


**Fig. S1. Inferring the homology of song components between *D. yakuba*, *D. teissieri*, and their male hybrids.**

**(a)** Representative song traces showing the two song types in *D. yakuba* (yPulse and yClack), *D. teissieri* (tSine and tPulse), and the two chimeric song types in the F1 hybrid males. The first chimeric song type appears to be a pulse song (hPulse) with an inter-pulse interval (hIPI) resembling tSine. The second hybrid chimeric song type appears to be clack song (hClack) followed by a tail resembling a truncated train of tPulse. **(b)** Randomly selected examples of song components defined in panel **a**. Notably, hIPI shares the most similar waveforms with tSine. **(c)** Histogram of carrier frequencies for the defined song components. Numbers of total events and fly individuals (in parentheses) are noted on the right. Again, hIPI shares the most similar carrier frequencies with tSine. Based on the comparison of waveforms and carrier frequencies, we inferred the following homologous relationships among song components: yPulse is homologous to hPulse; tSine to hIPI, yClack to hClack, tPulse to hCT.

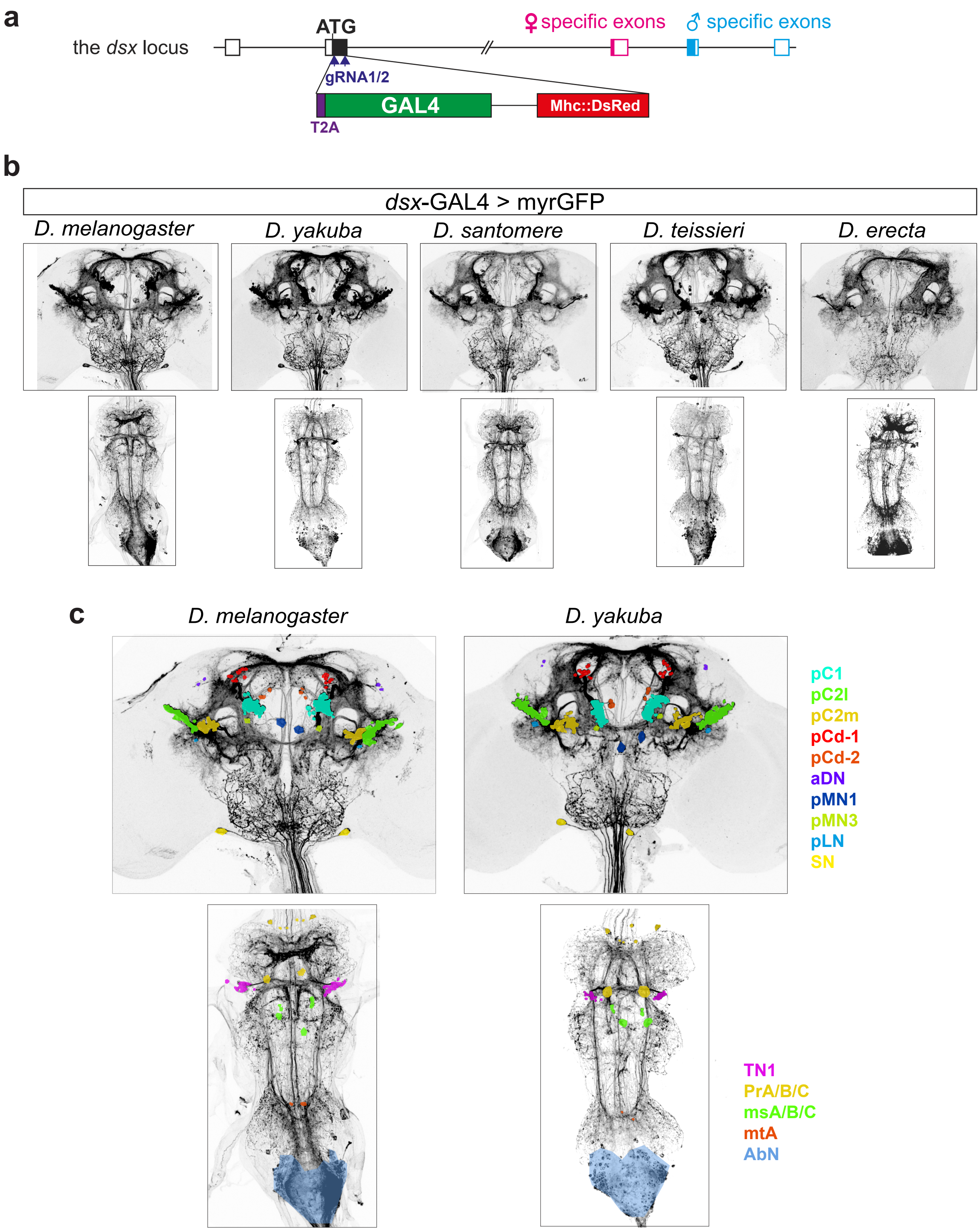


**Fig. S2. Generation of *dsx*-GAL4 knock-in transgenes.**

**(a)** CRISPR/Cas9-mediated targeting strategy of generating *dsx*-GAL4 alleles in *D. melanogaster*, *D. yakuba*, *D. santomea*, *D. teissieri*, and *D. erecta*. **(b)** Representative confocal images of *dsx*-GAL4>myrGFP in each species. **(c)** Confocal images of *D. melanogaster* and *D. yakuba* highlighting the cell bodies of previously defined *dsx*+ clusters[39,40]in different colors. All *dsx*+ clusters can be unambiguously defined in both species based on the location of cell bodies and arborization patterns.

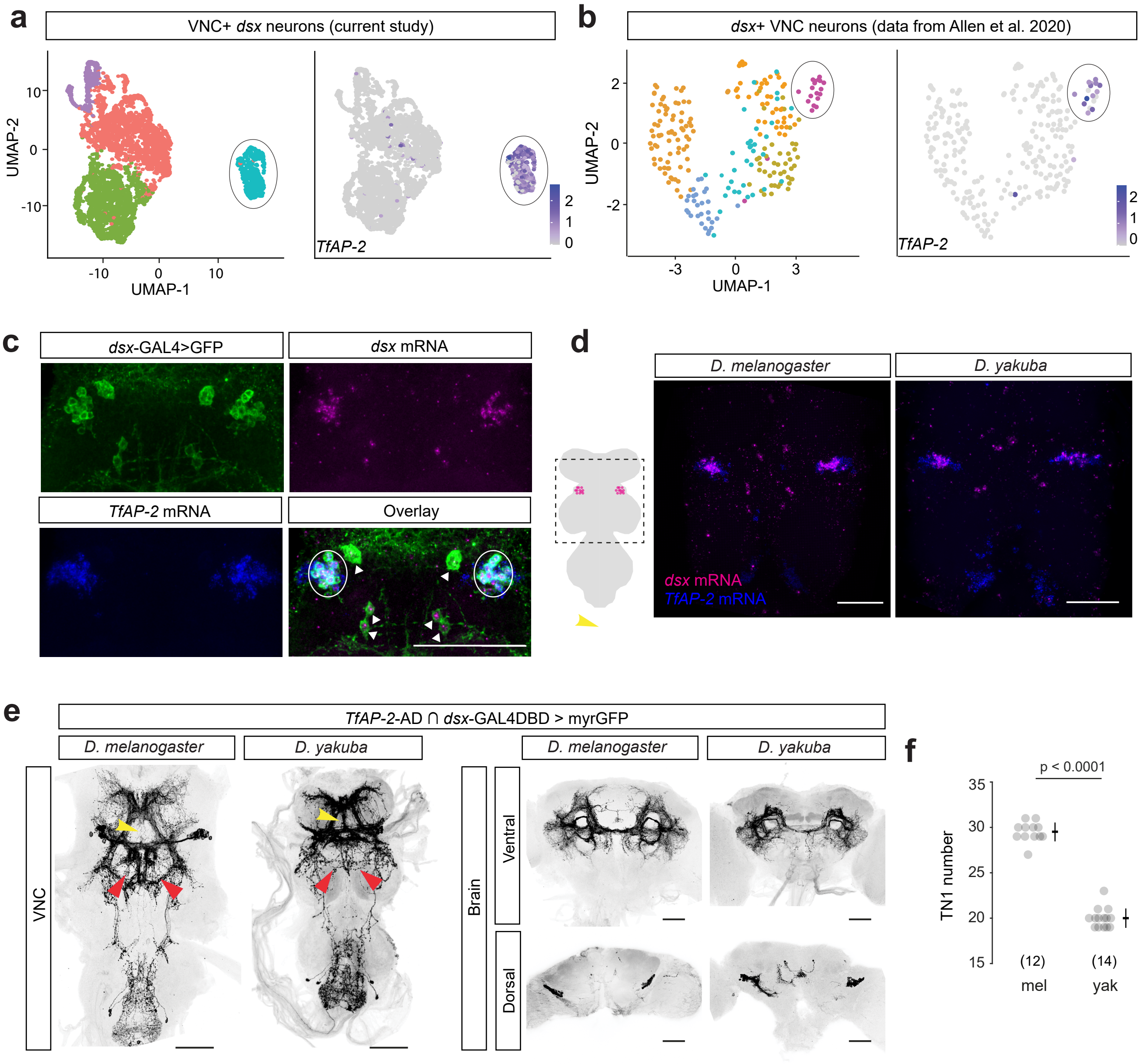


**Fig. S3. Identification of a conserved molecular marker for TN1 neurons in *D. melanogaster* and *D. yakuba*.**

**(a)** UMAP representations of molecular cell types in *dsx*+ neurons that express VNC markers (left) and the corresponding gene expression of *TfAP-2* (right). The circled cluster represents the putative TN1 neurons. **(b)** UMAP representations of molecular cell types in *dsx*-expressing neurons extracted from the published VNC scRNAseq dataset[38](left) and the corresponding gene expression of *TfAP-2* (right) in *D. melanogaster*. The circled cluster represents the putative TN1 neurons. **(c)** Confocal stacks showing the co-expression of *dsx* and *TfAP-2* in TN1 neurons (cell bodied circled) and the absence of *TfAP-2* expression in non-TN1 *dsx* neurons (arrowhead) in the regions of accessory mesothoracic neuropil and mesothoracic neuromere. **(d)** Co-expression of *dsx* and *TfAP-2* in the TN1 region is conserved between *D. melanogaster* and *D. yakuba*. **(e)** In VNCs of both species, the genetic intersection *TfAP-2*-AD∩ *dsx*-GAL4DBD specifically labels TN1 neurons and reveals similar species differences in the posterior arbors (red arrows) and the anterior medial arbors (yellow arrows) as shown in **Fig. 2b**. This intersection also labels pC2l neurons in the brain. **(f)** The genetic intersection *TfAP-2*-AD∩*dsx*-GAL4DBD labels the complete set of TN1 neurons in both *D. melanogaster* (mel) and *D. yakuba* (yak). Data are presented as mean±SD with sample sizes shown in parentheses. We performed a one-way ANOVA. Scale bars: 50 uM.

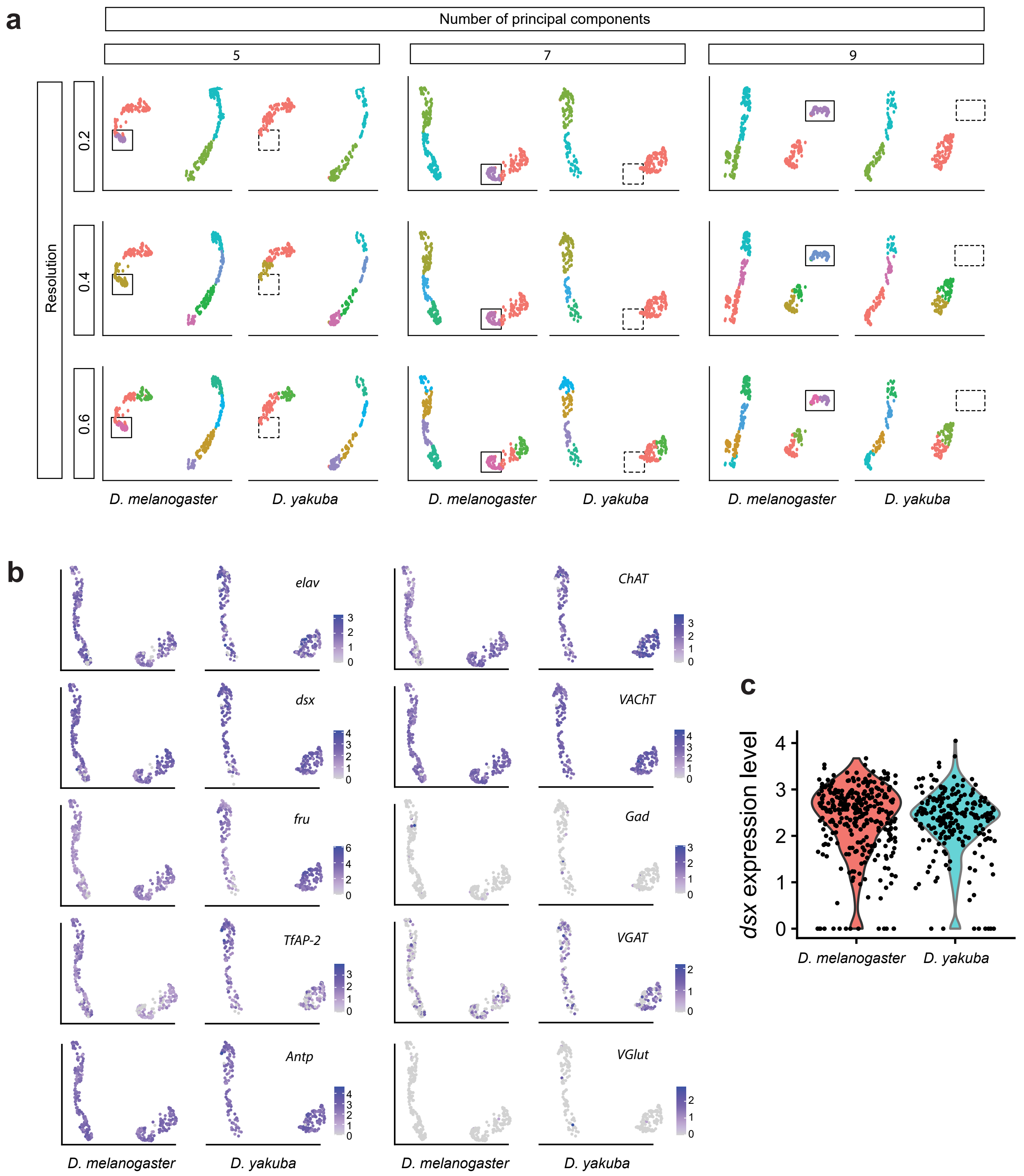


**Fig. S4. Species comparison of TN1 neurons in molecular subtypes and expression of key genes based on scRNAseq data.**

**(a)** UMAP representations highlighting the *D. melanogaster*-specific cluster across a range of principal components and Seurat "Find-Clusters" function resolutions in *D. melanogaster* and *D. yakuba*. **(b)** UMAP representations split by species showing the expression levels of *elav* (neuronal marker), *dsx*, *fru* (sex determination genes), *TfAP-2*, *Antp* (TN1 marker genes), *ChAT*, *VAChT* (cholinergic neuron markers), *Gad1*, *VGAT* (GABAergic neuron markers), and *VGlut* (glutamatergic neuron markers). **(c)** Violin plots showing the expression levels of *dsx* in TN1 neurons between *D. melanogaster* and *D. yakuba*. We performed a Mann-Whitney U test, adjusted p-value = 1. Each dot represents a single-cell transcriptome of a TN1 neuron.

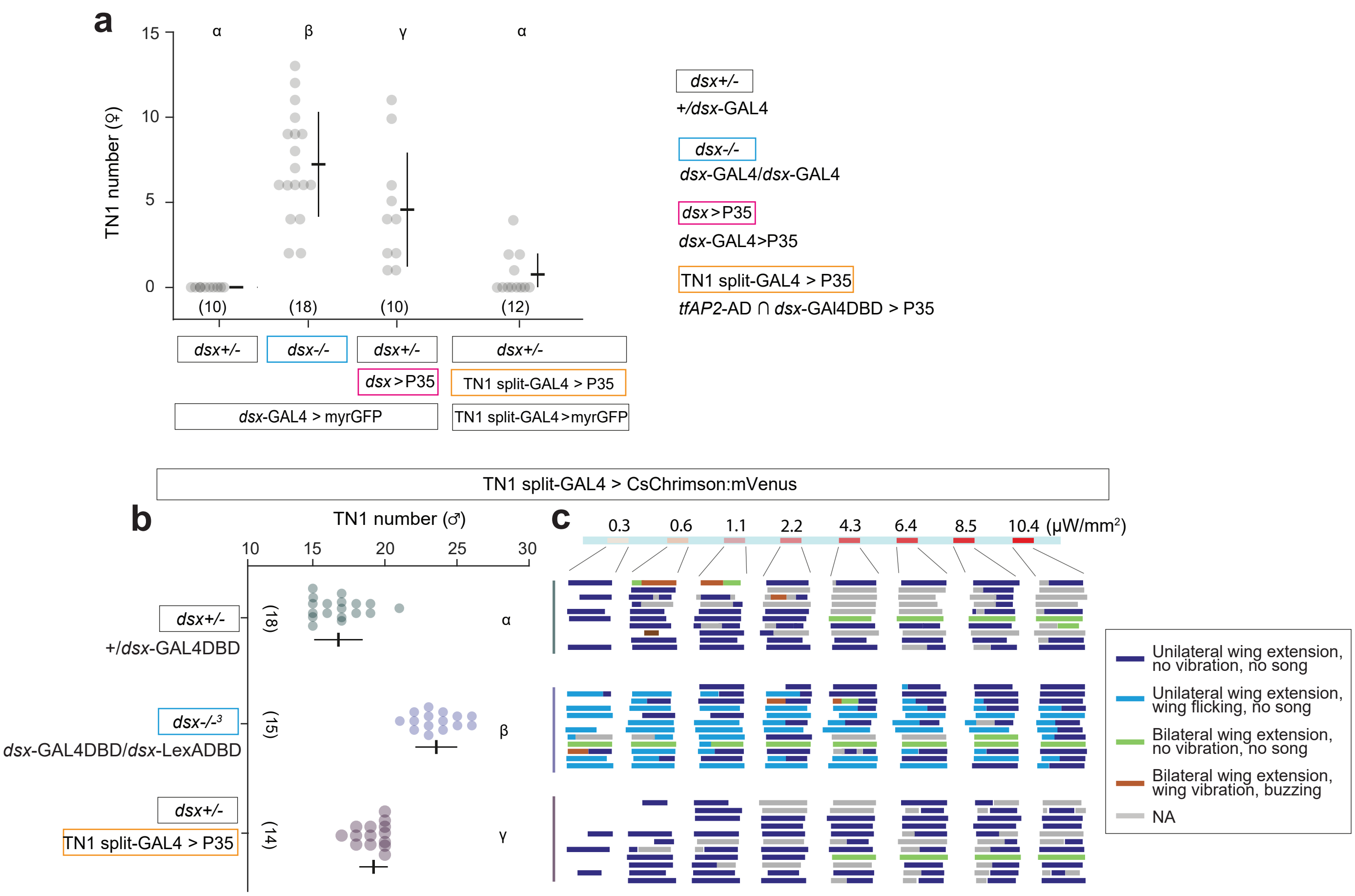


**Fig. S5. Additional data for TN1 resurrection in *D. yakuba*, related to Fig. 4.**

**(a)** P35 expression driven by the TN1-specific genetic intersection (TN1 split-GAL4>P35) in females had little effect in resurrecting TN1 neurons compared with *dsx* null (*dsx*-/-) and P35 expression driven by *dsx*-GAL4(*dsx*>P35). Genotype abbreviations are specified on the right. **(b)** Similarly, TN1 split-GAL4>P35 in males did not fully recapitulate the effects of *dsx* null (*dsx*-/-$^{3}$) or *dsx*-GAL4>P35 (see **Fig. 4a**). This suggests that the TN1 split-GAL4 may not drive P35 sufficiently high or early enough to fully block apoptosis as *dsx*-GAL4 did. **(c)** Behavioral responses of TN1 activation in decapitated males for the indicated genotypes. Unilateral wing extension coupled with wing flicking, albeit no song, was observed in *dsx* null males. Also see **Supplementary Video 2**. This result is consistent with its gain of TN1 neurons and the lack of *dsx* expression. The lack of a phenotype in TN1 split-GAL4>P35 flies is consistent with the insufficient resurrection of TN1 neurons. Data are presented as mean±SD with sample sizes shown in parentheses. We performed a Mann-Whitney U test with a Bonferroni Correction. Shared letters in panels **a**,**b** denote  no significant difference ($p > 0.05$).

**Supplementary Table 1. *Drosophila* genotypes used in this study.**

| Figure | Species | Male Genotypes |
|---|---|---|
| 1a-c,e; S3d | *D.melanogaster* | wildtype, Oregon |
| 1a-c; S1a-c; S3d | *D. yakuba* | wildtype, UCSD # 14021-0261.02 |
| 1a; | *D. santomea* | wildtype, UCSD # 14021-0271.00 |
| 1a,b; S1a-c | *D. teissieri* | wildtype, UCSD # 14021-0257.01 |
| 1a | *D. erecta* | wildtype, UCSD # 14021-0224.01 |
| 1b; S1a-c | *D. yakuba* x *D. teissieri* hybrid | NA |
| 1d,f,g; S2b, c; S3c | *D.melanogaster* | ; ; *dsx*-GAL4/UAS-myrGFP |
| 1d,f,g; S2b, c | *D. yakuba* | ; ; *dsx*-GAL4/UAS-myrGFP |
| 1d,f,g; S2b | *D. santomea* | ; UAS-myrGFP/+; *dsx*-GAL4/+ |
| 1d,f,g; S2b | *D. teissieri* | ; UAS-myrGFP/+; *dsx*-GAL4/+ |
| 1d,f,g; S2b | *D. erecta* | pBac{UAS-CsChrimson:tdTomato; 3XP3EYFP}-1A/+; ; *dsx*-GAL4/+ |
| 2a-c | *D.melanogaster* | w; 13H01-LexA/UAS-FRT-stop-FRT-myrGFP; LexAop-FLP/*dsx*-GAL4 |
| 2a-c | *D. yakuba* | w; 13H01-LexA (2180)/UAS-FRT-stop-FRT-myrGFP (2180); LexAop-FLP/*dsx*-GAL4 |
| 2d | *D.melanogaster* | w; 13H01-LexA/UAS-FRT-stop-FRT-CsChrimson::mVenus; LexAop-FLP/*dsx*-GAL4 |
| 2d | *D. yakuba* | w; 13H01-LexA (2180)/UAS-FRT-stop-FRT-CsChrimson::mVenus (2180); LexAop-FLP (2283)/*dsx*-GAL4 |
| 2e | *D.melanogaster* | w; 13H01-LexA/UAS-FRT-stop-FRT-Kir2.1::GFP (attP2); LexAop-FLP/*dsx*-GAL4 |
| 2e | *D. yakuba* | w; 13H01-LexA (2180)/UAS-FRT-stop-FRT-Kir2.1::GFP (2180); LexAop-FLP (2283)/*dsx*-GAL4 |
| 3a,b,f; S3a,b; S4a-c | *D.melanogaster* | ; pJFRC29-10XUAS-IVS-myr::GFP-p10 (attP40)/+; *dsx*-GAL4/pJFRC105-10XUAS-IVS-nlstdTomato (VK00040) |
| 3a,b,f; S4a-c | *D. yakuba* | ; ; *dsx*-GAL4/UAS-nls::tdTomato (2283) |
| 3c | *D.melanogaster* | w; VT017258-p65AD::Zp/UAS-myrGFP; $dsx^{Zp::Gdbd}$/+ |
| 3d | *D.melanogaster* | *w, 17D3*$^{GAL4}$/Y; UAS-FRT-stop-FRT-myrGFP/+; *dsx*-GAL4LexADBD,*TfAP-2*-AD/LexAop-FLP |
| 3e | *D.melanogaster* | *w, 17D3*$^{GAL4}$/Y; UAS-FRT-stop-FRT-Kir2.1::GFP (attP2)/+; *dsx*-GAL4LexADBD,*TfAP-2*-AD/LexAop-FLP |
| 4a | *D.melanogaster* | w; pJFRC29-10XUAS-IVS-myr::GFP-p10 (attP40)/+; *dsx*-GAL4/+ |
| 4a | *D.melanogaster* | w; pJFRC29-10XUAS-IVS-myr::GFP-p10 (attP40)/+; *dsx*-GAL4/dsx-GAL4 |
| 4a | *D.melanogaster* | w; pJFRC29-10XUAS-IVS-myr::GFP-p10 (attP40)/UAS-P35$^{BH1}$; *dsx*-GAL4/+ |
| 4a,b | *D. yakuba* | w, UAS-myrGFP (2180)/Y; ; *dsx*-GAL4/+ |
| 4a | *D. yakuba* | w, UAS-myrGFP (2180)/Y; ; *dsx*-GAL4/*dsx*-GAL4 |
| 4a,b | *D. yakuba* | w, UAS-myrGFP (2180)/Y; UAS-P35 (2180)/+; *dsx*-GAL4/+ |
| 4a | *D. yakuba* | w, UAS-myrGFP (2180)/Y; ; *dsx*-GAL4/*dsx*-GAL4LexADBD |
| 4a | *D. yakuba* | w, UAS-myrGFP (2180)/Y; UAS-P35 (2180)/+; *dsx*-GAL4/*dsx*-GAL4LexADBD |
| 4a | *D. santomea* | w; UAS-myrGFP/+; *dsx*-GAL4/+ |
| 4a | *D. santomea* | w; UAS-myrGFP/+; *dsx*-GAL4/*dsx*-GAL4 |
| 4c; S5b,c | *D. yakuba* | w; ; *TfAP-2*-AD, *dsx*-GAL4DBD, 20XUAS-CsChrimson-tdTomato (1730)/+ |
| 4c; S5b,c | *D. yakuba* | w; ; *TfAP-2*-AD, *dsx*-GAL4DBD, 20XUAS-CsChrimson-tdTomato (1730)/*dsx*-GAL4LexADBD |
| S5b,c | *D. yakuba* | w; UAS-P35 (2180)/+; *TfAP-2*-T2A-AD, *dsx*-GAL4DBD,20XUAS-CsChrimson-tdTomato (1730)/+ |
| S3e,f | *D.melanogaster* | w; pJFRC29-10XUAS-IVS-myr::GFP-p10 (attP40)/+; *TfAP-2*-AD, $dsx^{Zp::Gdbd}$/+ |
| S3e,f | *D. yakuba* | w, UAS-myrGFP (2180)/Y; ; *TfAP-2*-AD,*dsx*-GAL4DBD/+ |

| Figure | Species | Female Genotypes |
|---|---|---|
| S5a | *D. yakuba* | UAS-myrGFP/+; ; *dsx*-GAL4/+ |
| S5a | *D. yakuba* | UAS-myrGFP/+; ; *dsx*-GAL4/*dsx*-GAL4LexADBD |
| S5a | *D. yakuba* | UAS-myrGFP/+; UAS-P35 (2180)/+; *dsx*-GAL4/+ |
| S5a | *D. yakuba* | ; UAS-P35 (2180)/+; *TfAP-2*-AD, *dsx*-GAL4DBD, UAS-CsChrimson-tdTomato (1730)/+ |

**Supplementary Table 2. Primers to generate constructs for CRISPR/Cas9-mediated gene targeting.**

**Primers to generate the gRNA donor plasmids using pCFD4**

| Name | Sequences | gRNA sequences | Note |
|---|---|---|---|
| pCFD4_yakDsx_F | TATATAGGAAAGATATCCGGGTGAACTTCGGAGTCGATCATGTCCGAGTGTTTTAGAGCTAGAAATAGCAAG | GGAGTCGATCATGTCCGAGT | An extra G added to the 5' end of gRNA |
| pCFD4_yakDsx_R | ATTTTAACTTGCTATTTCTAGCTCTAAAACTCAATCGCAAGAACGGCGCCGACGTTAAATTGAAAATAGGTC | GGCGCCGTTCTTGCGATTGA | |
| pCFD4_sanDsx_F | same as pCFD4_yakDsx_F | GGAGTCGATCATGTCCGAGT | |
| pCFD4_sanDsx_R | same as pCFD4_yakDsx_R | GGCGCCGTTCTTGCGATTGA | |
| pCFD4_melDsx_F | same as pCFD4_yakDsx_F | GGAGTCGATCATGTCCGAGT | |
| pCFD4_melDsx_R | ATTTTAACTTGCTATTTCTAGCTCTAAAACTCAATCGAAAGAACGGCGCCGACGTTAAATTGAAAATAGGTC | GGCGCCGTTCTTTCGATTGA | |
| pCFD4_ereDsx_F | same as pCFD4_yakDsx_F | GGAGTCGATCATGTCCGAGT | |
| pCFD4_ereDsx_R | same as pCFD4_yakDsx_R | GGCGCCGTTCTTTCGATTGA | |
| pCFD4_teiDsx_F | same as pCFD4_yakDsx_F | GGAGTCGATCATGTCCGAGT | |
| pCFD4_teiDsx_R | ATTTTAACTTGCTATTTCTAGCTCTAAAACAGGGCACGTTGGCGCCGTTCGACGTTAAATTGAAAATAGGTC | GAACGGCGCCAACGTGCCCT | |
| pCFD4-yakTfAP2_F | TATATAGGAAAGATATCCGGGTGAACTTCGCTGCGTCCCGTGTGTCCGCGTTTTAGAGCTAGAAATAGCAAG | GCTGCGTCCCGTGTGTCCGC | |
| pCFD4-yakTfAP2_R | ATTTTAACTTGCTATTTCTAGCTCTAAAACTGCACCACCAGTCGCTGCACGACGTTAAATTGAAAATAGGTC | GTGCAGCGACTGGTGGTGCA | An extra G added to the 5' end of gRNA |
| pCFD4-melTfAP2_F | TATATAGGAAAGATATCCGGGTGAACTTCGCTGCGTCCCGTGTGGCCGCGTTTTAGAGCTAGAAATAGCAAG | GCTGCGTCCCGTGTGGCCGC | |
| pCFD4-melTfAP2_R | same as pCFD4-yakTfAP2_R | GTGCAGCGACTGGTGGTGCA | An extra G added to the 5' end of gRNA |

**Primer to amplify the homology arms**

| Name | Sequences | Sequences excluding the overlapping tips for Gibson assembly |
|---|---|---|
| yakDsx_Larm_F | AAAAGTGCCACCTGACGTCTTGGAAGCAAGTGTTTTGTGTGA | TGGAAGCAAGTGTTTTGTGTGA |
| yakDsx_Larm_R | AGCAGCGATCCGCGACCCTCGGACATCGTGTCGCTATTCCA | GGACATCGTGTCGCTATTCCA |
| yakDsx_Rarm_F | GTTGTGGTTTGTCCAAACTCCCAACGTGCCCTTGGGTAAG | CCAACGTGCCCTTGGGTAAG |
| yakDsx_Rarm_R | GCCTTTTTACGGTTCCTGGCATTGTCATTCCCCCTGGCTG | ATTGTCATTCCCCCTGGCTG |
| sanDsx_Larm_F | AAAAGTGCCACCTGACGTCTCGAAATAATCCTGCAGTTAAATATG | CGAAATAATCCTGCAGTTAAATATG |
| sanDsx_Larm_R | same as yakDsx_Larm_R | GGACATCGTGTCGCTATTCCA |
| sanDsx_Rarm_F | same as yakDsx_Rarm_F | CCAACGTGCCCTTGGGTAAG |
| sanDsx_Rarm_R | yakDsx_Rarm_R | ATTGTCATTCCCCCTGGCTG |
| melDsx_Larm_F | AAAAGTGCCACCTGACGTCTAGTAATCCGGCATATTCTTACCCAT | AGTAATCCGGCATATTCTTACCCAT |
| melDsx_Larm_R | same as yakDsx_Larm_R | GGACATCGTGTCGCTATTCCA |
| melDsx_Rarm_F | GTTGTGGTTTGTCCAAACTCCCAATGTGCCCTTGGGTAAG | CCAATGTGCCCTTGGGTAAG |
| melDsx_Rarm_R | GCCTTTTTACGGTTCCTGGCATTGTCATTCCCGCTGGCTG | ATTGTCATTCCCGCTGGCTG |
| teiDsx_Larm_F | AAAAGTGCCACCTGACGTCTCGAAAAAACCTTTCAGTTAAATTTG | CGAAAAAACCTTTCAGTTAAATTTG |
| teiDsx_Larm_R | same as yakDsx_Larm_R | GGACATCGTGTCGCTATTCCA |
| teiDsx_Rarm_F | GTTGTGGTTTGTCCAAACTCCTTGGGTAAGTGAATACCAT | CTTGGGTAAGTGAATACCAT |
| teiDsx_Rarm_R | GCCTTTTTACGGTTCCTGGCATTGTCATTCCCGCTGGCTG | ATTGTCATTCCCGCTGGCTG |
| ereDsx_Larm_F | AAAAGTGCCACCTGACGTCTAGTAATCCGGAATATTCTTACCCAT | AGTAATCCGGAATATTCTTACCCAT |
| ereDsx_Larm_R | same as yakDsx_Larm_R | GGACATCGTGTCGCTATTCCA |
| ereDsx_Rarm_F | same as melDsx_Rarm_F | CCAATGTGCCCTTGGGTAAG |
| ereDsx_Rarm_R | GCCTTTTTACGGTTCCTGGCTGGACGGGGCCAGGATGAAA | TGGACGGGGCCAGGATGAAA |

| | | |
|---|---|---|
| yakTfAP2_Larm_F | AAAAGTGCCACCTGACGTCTCTGGCGCGC GACTAGTTCTC | CTGGCGCGCGACTAGTTCTC |
| yakTfAP2_Larm_R | AGCAGCGATCCGCGACCCTCGCCGGAATG CGTTGTGTAC | CCGGAATGCGTTGTGTAC |
| yakTfAP2_Rarm_F | GTTGTGGTTTGTCCAAACTCCACCAGTCGC TGCAGTCCG | CACCAGTCGCTGCAGTCCG |
| yakTfAP2_Rarm_R | GCCTTTTTACGGTTCCTGGCTGTGTGTGAG AGTCCTTTGGG | TGTGTGTGAGAGTCCTTTGGG |
| melTfAP2_Larm_F | AAAAGTGCCACCTGACGTCTCTGGCGAGC GACTAGTTCTC | CTGGCGAGCGACTAGTTCTC |
| melTfAP2_Larm_R | same as yakTfAP2_Larm_R | CCGGAATGCGTTGTGTAC |
| melTfAP2_Rarm_F | same as yakTfAP2_Rarm_F | CACCAGTCGCTGCAGTCCG |
| melTfAP2_Rarm_R | same as yakTfAP2_Rarm_R | TGTGTGTGAGAGTCCTTTGGG |

**Supplementary Table 3. Sources of Drosophila lines in this study.**

| Fly lines | Source | Injected lines | Injected plasmids |
|---|---|---|---|
| *D. melanogaster* , ; ; *dsx* -GAL4/+ | This study | *D. melanogaster* Oregon | *mel-dsx* -T2A-GAL4, Mhc-DsRed, described in Methods |
| *D. yakuba* , ; ; *dsx* -GAL4/+ | This study | UCSD # 14021-0261.02 | *yak-dsx* -T2A-GAL4, Mhc-DsRed, described in Methods |
| *D. santomea* , ; ; *dsx* -GAL4/+ | This study | UCSD # 14021-0271.00 | *san-dsx* -T2A-GAL4, Mhc-DsRed, described in Methods |
| *D. teissieri* , ; ; *dsx* -GAL4/+ | This study | UCSD # 14021-0257.01 | *tei-dsx* -T2A-GAL4, Mhc-DsRed, described in Methods |
| *D. erecta* , ; ; *dsx* -GAL4/+ | This study | UCSD # 14021-0224.01 | *ere-dsx* -T2A-GAL4, Mhc-DsRed, described in Methods |
| *D. melanogaster* , ; ; *TfAP-2* -GAL4/+ | This study | *D. melanogaster* Oregon | mel-*TfAP-2* -T2A-GAL4, Mhc-DsRed, described in Methods |
| *D. melanogaster* , ; ; *TfAP-2-* AD/+ | This study | *D. melanogaster* Oregon | mel-*TfAP-2* -T2A-AD, Mhc-DsRed, described in Methods |
| *D. yakuba* , ; ; *dsx* -GAL4DBD/+ | This study | UCSD # 14021-0261.02 | *yak-dsx* -T2A-GAL4DBD, Mhc-DsRed, described in Methods |
| *D. yakuba* , ; ; *dsx* -GAL4LexADBD/+ | This study | UCSD # 14021-0261.02 | *yak-dsx* -T2A-LexADBD, Mhc-DsRed, described in Methods |
| *D. yakuba* , ; ; *TfAP-2* -AD/+ | This study | UCSD # 14021-0261.02 | *yak* -*TfAP-2* -T2A-AD, Mhc-DsRed, described in Methods |
| *D. yakuba* , w; UAS-P35 (2180); | This study | *D. yakuba* 2180 (mapped to 2 chr) | 10XUAS-P35, described in Methods |
| *D. yakuba* , w; 13H01-LexA (2180); | This study | *D. yakuba* 2180 (mapped to 2 chr) | 13H01-LexA |
| *D. yakuba* , w; ;LexAop-FLP (2283) | This study | *D. yakuba* 2283 | pJFRC79-8XLexAop2-FLPL |
| *D. yakuba* , w; UAS-FRT-stop-FRT-myrGFP (2180); | This study | *D. yakuba* 2180 (mapped to 2 chr) | pJFRC177-10XUAS-FRT>-dSTOP-FRT>-myr::GFP |
| *D. yakuba* , w; UAS-FRT- stop-FRT-CsChrimson::mVenus (2180); | This study | *D. yakuba* 2180 (mapped to 2 chr) | UAS- FRT-stop-FRT-CsChrimson::mVenus |
| *D. yakuba* , w; UAS- FRT-stop-FRT-Kir2.1::GFP (2180); | This study | *D. yakuba* 2180 (mapped to 2 chr) | pJFRC56-UAS- FRT-stop-FRT-Kir2.1::GFP |
| *D. teissieri* , w; UAS-myrGFP (2180); | This study | *D. teissieri* w- | pBac{UAS-myrGFP; w+} |
| *D. yakuba* , w; ; UAS-nls::tdTomato (2283) | from David Stern's lab | *D. yakuba* 2283 | pJFRC105-10XUAS-IVS-nlstdTomato |
| *D. yakuba* , w; UAS-myrGFP (2180); | from David Stern's lab | *D. yakuba* 2180 (mapped to X chr) | pJFRC29-10XUAS-IVS-myr::GFP-p10 |
| *D. erecta* , pBac{UAS-CsChrimson:tdTomato; 3XP3EYFP}-1A; ; | from David Stern's lab | UCSD # 14021-0224.01 | pBac{UAS-CsChrimson:tdTomato; 3Xp3EYFP} |
| *D. melanogaster* , w; ; UAS-myrGFP | Janelia | NA | NA |
| *D. melanogaster* , w; pJFRC29-10XUAS-IVS-myr::GFP-p10 (attP40); pJFRC105-10XUAS-IVS-nlstdTomato (VK00040) | Janelia | NA | NA |
| *D. yakuba* , w; ; UAS-myrGFP | Ding et al. 2019 | NA | NA |
| *D. santomea* , w; UAS-myrGFP; | Ding et al. 2019 | NA | NA |
| *D. yakuba* , w; ; 20XUAS-CsChrimson-tdTomato (1730) | Ding et al. 2019 | NA | NA |
| *D. melanogaster* , w; 13H01-LexA; | Shirangi et al. 2016 | NA | NA |
| *D. melanogaster* , w; ; *dsx* $^{Zp::Gdbd}$ | Shirangi et al. 2016 | NA | NA |
| *D. melanogaster* , w; pJFRC56-10XUAS-FRT>STOP>Kir2.1::GFP (attP2); | Shirangi et al. 2016 | NA | NA |
| *D. melanogaster* , w; VT017258-*p65* AD::Zp; | Shirangi et al. 2016 | NA | NA |
| *D. melanogaster* , w,*17D3* $^{GAL4}$; ; | Wu et al. 2019 | NA | NA |
| *D. melanogaster* , w; UAS-P35 $^{BH1}$; | BDSC # 5072 | NA | NA |
| *D. melanogaster* , wildtype | Oregon | NA | NA |
| *D. yakuba* , wildtype | UCSD # 14021-0261.02 | NA | NA |
| *D. santomea* , wildtype | UCSD # 14021-0271.00 | NA | NA |
| *D. teissieri* , wildtype | UCSD # 14021-0257.01 | NA | NA |
| *D. erecta* , wildtype | UCSD # 14021-0224.01 | NA | NA |

**Supplementary Table 4. Sources of antibodies, probes, and hairpins used in IHC and HCR.**

| Primary antibodies | Source | Concentration |
|---|---|---|
| Rat-anti-Elav | DHSB 7E8A10 | 1:200 |
| Mouse-anti-Brp | DSHB nc82 | 1:30 |
| Rabbit-anti-DsxM | Han et al. PNAS 2022 | 1:500 |
| Chicken-anti-GFP | Abcam #ab13970 | 1:600 |
| Rabbit-anti-DsRed | TaKaRa #632496 | 1:500 |

| Secondary antibodies | Source | Concentration |
|---|---|---|
| Goat-anti-Chicken Alexa Fluor 488 | Thermo Fisher A-11039 | 1:500 |
| Goat-anti-Mouse Alexa Fluor 568 | Thermo Fisher A-11031 | 1:500 |
| Goat-anti-Rabbit Alexa Fluor 568 | Thermo Fisher A-11011 | 1:500 |
| Goat-anti-Rat Alexa Fluor 647 | Thermo Fisher A-21247 | 1:500 |
| Goat-anti-Mouse Alexa Fluor 647 | Thermo Fisher A-28181 | 1:500 |

| RNA probes | Source | Concentration |
|---|---|---|
| mel-*dsx* -B1 | Molecular Instruments | 16 nM |
| yak-*dsx* -B1 | Molecular Instruments | 16 nM |
| mel-*17D3* -B2 | Molecular Instruments | 16 nM |
| yak-*17D3* -B2 | Molecular Instruments | 16 nM |
| mel-*TfAP-2* -B3 | Molecular Instruments | 16 nM |
| yak-*TfAP-2* -B3 | Molecular Instruments | 16 nM |

| RNA hairpin amplifiers | Source | Concentration |
|---|---|---|
| B1-Alexa Fluor 488 | Molecular Instruments | 30 pmol (15 pmol/complementary hairpin) |
| B1-Alexa Fluor 647 | Molecular Instruments | 30 pmol (15 pmol/complementary hairpin) |
| B2-Alexa Fluor 647 | Molecular Instruments | 30 pmol (15 pmol/complementary hairpin) |
| B3-Alexa Fluor 546 | Molecular Instruments | 30 pmol (15 pmol/complementary hairpin) |
